# A TNL receptor mediates microbiome feedbacks in Arabidopsis

**DOI:** 10.1101/2025.02.25.640125

**Authors:** Henry Janse van Rensburg, Niklas Schandry, Jan Waelchli, Katja Stengele, Selma Cadot, Katharina Jandrasits, Hiroaki Adachi, Claude Becker, Klaus Schlaeppi

**Author notes:** Contributed equally (Henry Janse van Rensburg, Niklas Schandry). Corresponding authors: Niklas Schandry Claude Becker Klaus Schlaeppi.

## Abstract

Plant performance depends on the soil microbiome. While microbiome feedbacks are well documented, the mechanisms by which plants perceive and mediate these feedbacks remain unclear. We established a framework using two distinct microbiomes in the same soil, where one led to enhanced growth of the *Arabidopsis thaliana* accession Col-0. Screening 410 accessions revealed substantial variation in growth feedbacks, which we used for genome-wide association mapping. We identified the immune receptor *Mediator of Microbiome Feedback 1* (*MMF1*) as a candidate gene involved in microbiota feedbacks. Characterisation in the reference accession Col-0 revealed that *mmf1* mutants lost the beneficial growth feedback, had an altered root bacterial community, and failed to induce a defence-related transcriptional response observed in wild-type plants. The discovery of MMF1 implies that integration of microbial signals optimises host microbiome composition and immune status to enhance growth.

## Main

Microbiomes are widely recognized for supporting their hosts with many vital services, often leading to improved host performance. Similar to the gut microbiome of humans, plant root microbiomes contribute to nutrition homeostasis, stress tolerance and host health (*1*). For plants, above-ground performance depends to a large extent on below-ground bacteria, fungi and other microbes that collectively function as a soil microbiome (*1*, *2*). Soil microbiomes consist of functionally diverse members providing beneficial services to plants, including nutrient acquisition, growth promotion, stress tolerance, and suppression of plant pathogens, but they can also contain pathogenic members that reduce plant fitness (*3*, *4*). Plant responses, which are causally linked to a soil microbiome, are referred to as microbiome feedbacks (*5*). This term is akin to plant-soil feedbacks describing the ecology of soil properties (more broadly, but including the microbiome) that influence plant performance (*6*, *7*). Microbiome feedbacks are highly relevant in agricultural contexts, as soil microbiome composition has been linked to agronomic traits of crops such as maize and potato (*8*, *9*), to field productivity (*10*), and to over- and under-yielding areas in fields (*11*, *12*). While different soil microbiomes have often been seen as drivers of differential plant responses (*13*, *14*), little is known about the mechanisms by which plants perceive complex soil microbiomes and mediate microbiome feedbacks.

Plants have evolved a sophisticated immune system that regulates interactions with beneficial microbes while defending against pathogens (*15–17*). Plant immunity operates through two key pathways: pattern-triggered immunity (PTI) and effector-triggered immunity (ETI). PTI is initiated at the cell surface by pattern-recognition receptors that recognize microbe-associated molecular patterns and provides basal defence to deter non-adapted microbes. In contrast, ETI is activated within the cells by nucleotide-binding leucine-rich repeat receptors (NLRs), which detect pathogen effectors or their activities, leading to definite immune responses (*18–20*). NLRs are divided into Toll/interleukin-1 receptor (TIR-NLRs; TNLs) and coiled-coil NLRs (CNLs). *Arabidopsis thaliana* Columbia-0 (Col-0) (hereafter: Arabidopsis) encodes 54 TNLs and 88 CNLs, highlighting the complexity of the plant immune system (*21*). Both PTI and ETI rely on signalling molecules such as salicylic acid (SA) and reactive oxygen species (ROS) (*19*, *20*). Although these pathways were primarily studied in the context of binary interactions with pathogenic or beneficial microbes, recent evidence suggests that PTI and ETI, and signalling via SA and ROS also influence plant-associated microbiomes. These immune pathways shape microbiome composition, modulating microbial interactions and contributing to overall plant health and resilience (*16*, *22–24*).

To study microbiome feedbacks, we developed an experimental framework based on two distinct soil microbiomes in the same soil type. These soil microbiomes are generated by growing wild-type maize, which produces benzoxazinoids (BXs), or BX-deficient mutant maize. BXs are bioactive compounds of *Poaceae* (*25*) that, among multiple functions, shape the root and rhizosphere microbiomes of plants (*14*, *26*, *27*). After three months of growth, the wild-type and mutant plants are removed, leaving behind BX-conditioned (BX_plus_) and its control (BX_minus_) soil microbiomes, respectively. We have described growth and defence feedbacks of maize and wheat in response to BX_minus_ and BX_plus_ soil microbiomes (*14*, *28*), and we have proven agronomic relevance of these microbiome feedbacks (*28*, *29*). Recently, we reported that such microbiome feedbacks also exist in the model plant Arabidopsis (*30*). The reference accession Col-0 responded with improved growth and enhanced defence when grown on the BX_plus_ compared to the BX_minus_ soil microbiome. As a plausible mechanistic explanation for this apparent lack of a growth-defence trade-off, the plants growing on the BX_plus_ soil microbiome were primed for SA defences (*31*). Through sterilization and microbiome complementation experiments, we demonstrated that the feedbacks were BX-dependent, but that the soil microbiomes drove the differential plant responses (*14*, *28*, *29*).

Microbiome feedbacks often differ between different genotypes of response plant (*32*, *33*). Also, in our work with maize and wheat responding to BX_minus_ and BX_plus_ soil microbiomes, we had observed a genetic component in microbiome feedbacks (*14*, *28*), but the underlying genetic basis for such feedbacks to soil microbiomes remains unknown. Here, we aimed to uncover the genetic basis underlying the microbiome feedbacks in Arabidopsis Col-0. To this end, we harnessed the genetic diversity among Arabidopsis accessions to study their growth feedbacks and used high-throughput phenotyping in combination with genome-wide association (GWA) mapping (**Extended Data Fig. 1**). We identified and functionally validated a yet uncharacterized TNL driving differential plant growth on the two soil microbiomes in the reference accession.

## Results

### Arabidopsis displays substantial phenotypic variation in microbiome feedbacks

Col-0 responded to the BX_plus_ soil microbiome with larger rosettes compared to the BX_minus_ soil microbiome, confirming earlier work (**Fig. 1a**, (*30*)). The effect size of the growth feedback was less pronounced at earlier timepoints, and a linear-mixed-effects model revealed a positive *time* × *microbiome* interaction, indicating that BX_plus_ plants grow faster than BX_minus_ plants over time (**Fig. 1b**). We set out to identify candidate loci that confer this growth feedback in this accession.

**Figure 1.**
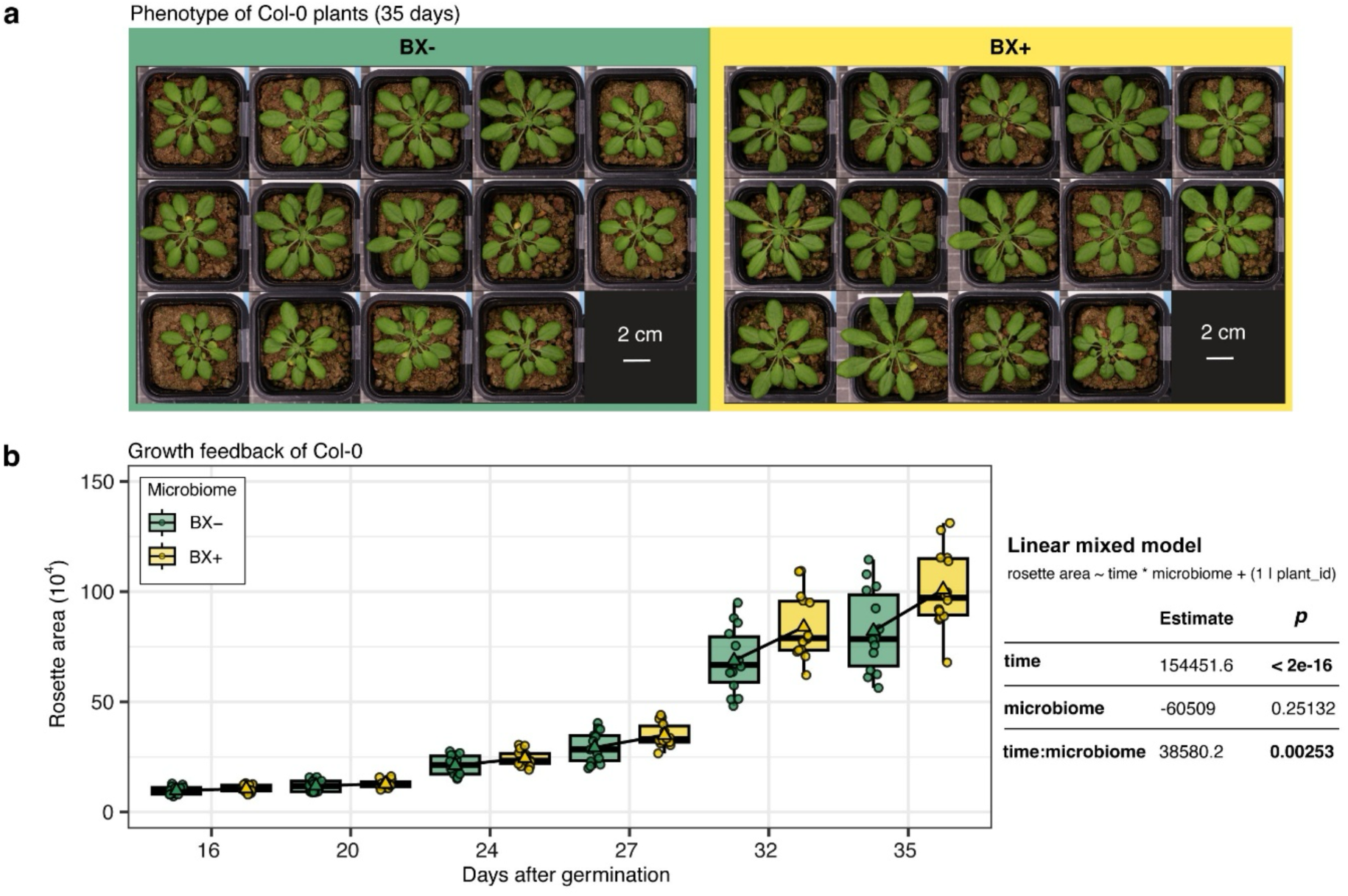
Growth feedbacks of Col-0 plants in response to the two soil microbiomes. (**a**) Phenotypes of Arabidopsis Col-0 plants grown on the BXminus and BXplus soil microbiomes at 35 days after germination. (**b**) Time course growth feedbacks of Arabidopsis Col-0 plants grown on the BXplus compared to the BXminus microbiome (points = individual plants). Linear-mixed-effects model of Col-0 rosette area across *time* and *microbiome* (minus vs plus).

We grew 410 natural, genome-sequenced Arabidopsis accessions (*34*) in both control (BX_minus_) and BX-conditioned (BX_plus_) soil microbiomes in a climate chamber equipped with an automated top-view imaging system. The process from imaging to determination of rosette area and feedback calculation is illustrated with the two contrasting accessions IP-Hue-3 and Hadd-3, which grew better and worse on the BX_plus_ soil microbiome, respectively (**Fig. 2a**). Plants were imaged twice a day, and plant morphometric traits were extracted using ARADEEPOPSIS (*35*). We focussed on the first 20 days after germination (DAG) to avoid confounding phenotyping effects such as early flowering and overlapping leaves. We used rosette area as a measurement of plant size to calculate the daily growth feedback as the difference in rosette area between BX_plus_ and BX_minus_ microbiomes, i.e., the accessions IP-Hue-3 and Hadd-3 exhibited positive and negative feedbacks, respectively (**Fig. 2b**).

**Figure 2.**
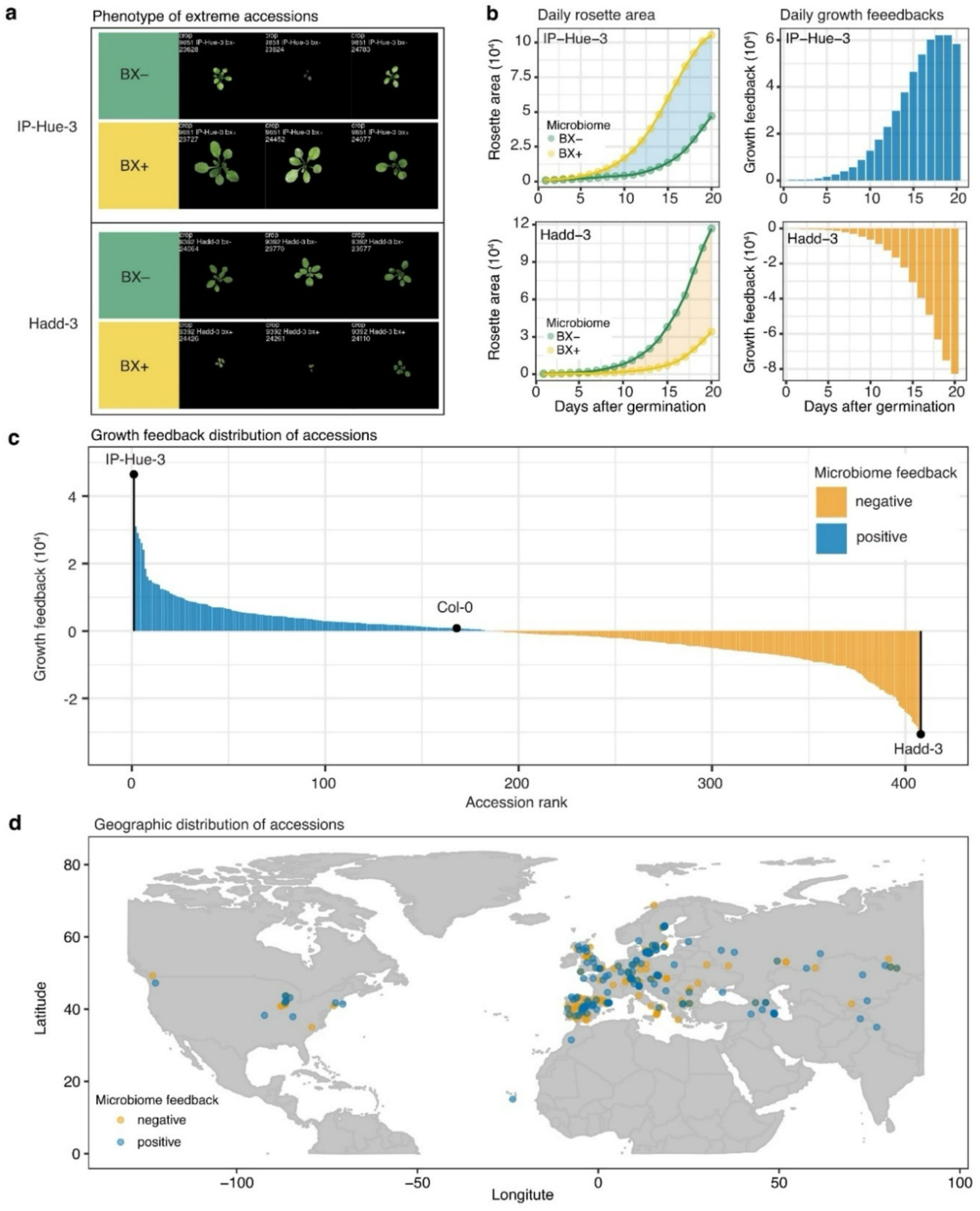
Natural variation in growth feedbacks among Arabidopsis accessions. (**a**) Growth of the two Arabidopsis accessions (IP-Hue-3 and Hadd-3) that exhibited the largest opposing growth feedbacks in response to the BXminus and BXplus microbiome at the conclusion of the experiment (20 days after germination (DAG)). (**b**) The daily rosette area measured in pixels extracted by ARADEEPOPSIS (left), growth feedback (BXplus – BXminus) (right) for the first 20 DAGs for IP-Hue-3 and Hadd-3 grown on the two soil microbiomes. Shaded areas (left) and coloured bars (right) indicate positive (blue, IP-Hue-3) or negative (orange, Hadd-3) microbiome feedbacks in response to the BXplus microbiome. (**c**) Microbiome feedback of the 410 *A. thaliana* accessions used in the genome-wide association study at the conclusion (DAG 20) of the experiment. The position of IP-Hue-3, Hadd-3 and Col-0 are indicated on the graph. Data is from two phenotyping experiments (experiment I and II, **Table S1**), each consisting of a subset of 210 accessions (3 replicates per accessions per soil), with 10 overlapping accessions present in each of the experiments. (**d**) The map shows the geographic distribution of these accessions, coloured by their microbiome feedback (blue for positive; orange for negative).

Screening of natural accessions revealed substantial variation in growth feedbacks to BX_plus_ and BX_minus_ microbiomes (**Fig. 2c**, **Dataset 1** and **Suppl. File 1**). Consistent with the above, Col-0 expressed a subtle positive feedback at the early timepoint of the GWA experiment. Across the 410 accessions, 48% (196 accessions) displayed positive feedbacks, while 52% (214 accessions) showed negative feedbacks. At the two extremes of the distribution, the two soil microbiomes accounted for ∼2-fold (IP-Hue-3) and >3-fold (Hadd-3) differences in rosette area in response to the two microbiomes. The observed feedback responses were independent of the accessions’ geographical origin (**Fig. 2d**, **Extended Data Fig. 2a**).

Since BXs are allelopathic to plants (*36*, *37*), we tested whether the accessions’ growth differences could be due to varying sensitivity to residual BXs in soil. BXs differ between the two soil variants, with MBOA as the most abundant BX compound (*29*, *38*). We screened a subset of accessions with the strongest opposing growth feedbacks for sensitivity to MBOA. All accessions exhibited shorter roots in the presence of MBOA (**Extended Data Fig. 3a**, **Dataset 2** and **Suppl. File 2**), which is indicative of phytotoxicity. However, there was no correlation between the microbiome response of the accessions and their sensitivity to MBOA (**Extended Data Fig. 3b**). These results indicated that the observed growth feedbacks are not linked to varying sensitivity to the BXs present in the soil.

In conclusion, we found substantial phenotypic variation in growth feedbacks of Arabidopsis accessions to two soil microbiomes, providing a powerful resource for association studies on the genetics that drive the differences in feedbacks.

### Candidate gene loci are involved in plant growth and defence

We performed a genome-wide association study (GWAS) using the phenotypic variation in growth feedbacks of the 410 accessions (**Fig. 2c**) and employed a linear mixed model, accounting for population structure (*39*). Noting temporal variation in daily growth feedbacks (**Extended Data Fig. 2b**), we ran separate GWAS on daily data and calculated associations between single nucleotide polymorphisms (SNPs) and the phenotypes for each of the 20 DAGs (**Dataset 3**).

Across daily GWAS runs, we identified three genomic regions with significant associations to the growth feedback (**Fig. 3a**, **Dataset 3** and **Suppl. File 3**). Genes in the regions of the three peaks were related to plant growth and plant defence. We then focused on SNPs that were significant for multiple consecutive days in the repeated GWAS (see Methods). The top-ranked SNP (chromosome 3, position 16203882) was in peak #2 (**Fig. 3a**), of which an alternative allele was found in 11 accessions (**Fig. 3b**, **c**). On average, these accessions showed better growth in response to the BX_plus_ microbiome compared to accessions with the reference allele. The SNP is in close proximity to the TNL receptor gene *AT3G44630* (position 16200156; 3.7kb upstream) and a more distal second TNL (*AT3G44670*; position 16218427; 14.5kb downstream; **Fig. 3d**). We focussed on the uncharacterized *AT3G44630* because of i) the closer physical proximity to the top-ranking SNP (**Fig. 3d**), and ii) the differential expression of *AT3G44630* in roots of Arabidopsis grown on BX_minus_ and BX_plus_ soils (*30*). In fact, from the 33 candidate genes identified from the top 100 GWA peaks, only *AT3G44630* was both in close proximity to a significant GWA peak (**Fig. 3a**) and displayed BX-dependent differential expression in the roots of Col-0 (**Fig. 3e**). We named this gene *Mediator of Microbiome Feedback 1* (*MMF1*). Structural predictions indicated high similarity to the well-characterized TNL, Recognition of Peronospora Parasitica 1 (RPP1, (*40*)), suggesting that MMF1 also functions as a tetrameric TNL resistosome (**Fig. S1**, **Suppl. Results**). However, unlike RPP1, MMF1 contains unique N- and C-terminal extensions. Taken together, GWAS identified a genomic locus on chromosome 3 that includes *MMF1* as a candidate gene that could explain the differential growth feedbacks of Arabidopsis accessions in response to the BX microbiomes.

**Figure 3.**
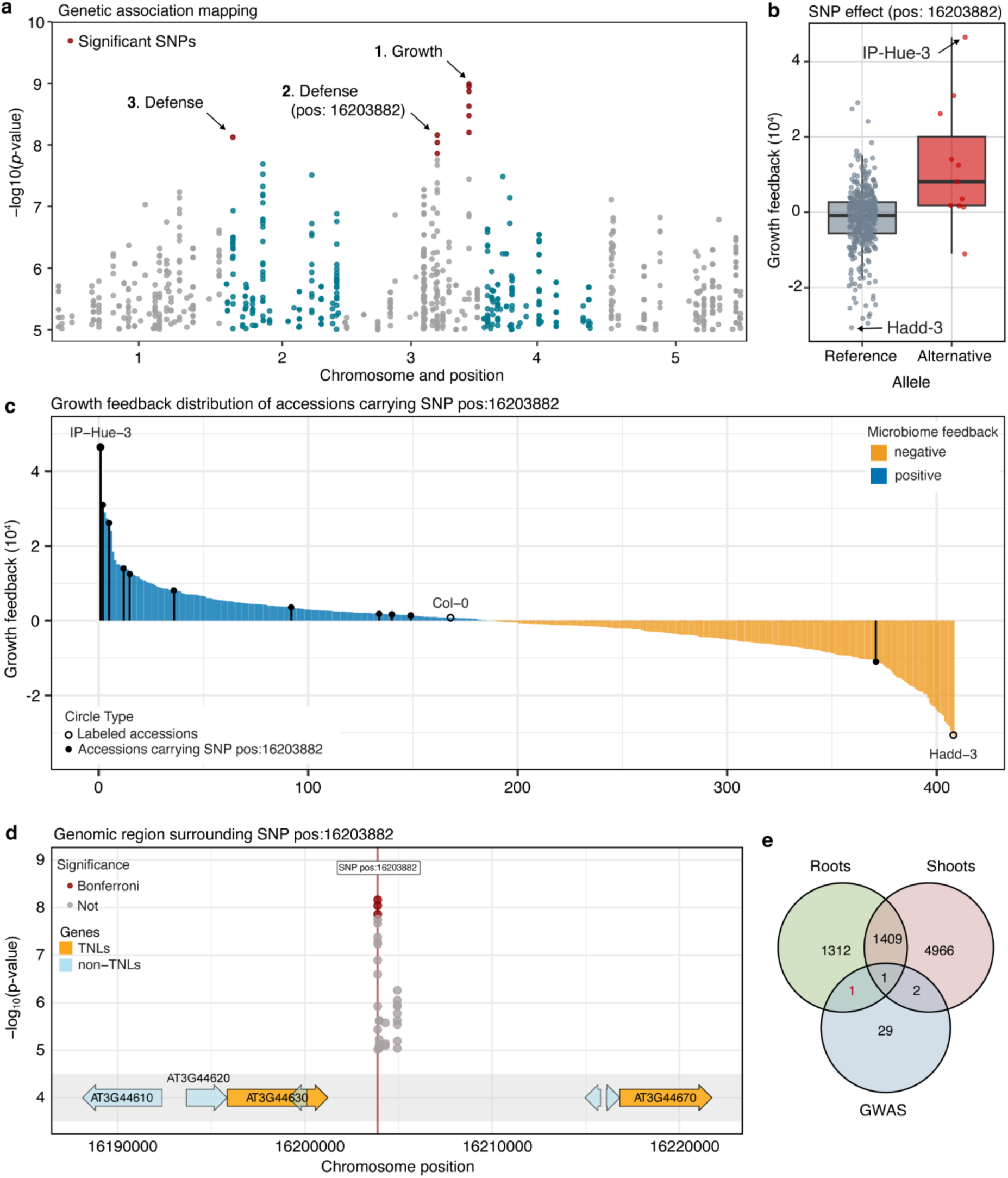
Genetic association mapping of growth feedbacks in response to the BXplus microbiome. (**a**) Manhattan plot showing the -log10-transformed *p*-values for the top 500 single nucleotide polymorphisms (SNPs) across the 5 chromosomes of Arabidopsis. SNPs that are significantly associated with growth feedbacks after Bonferroni correction are shown in red. The biological functions of genes adjacent to the three top-ranking SNPs are indicated. (**b**) Average growth feedback among accessions carrying the alternative allele of the SNP at position 16,203,882 (positive feedback) compared to those with the reference allele (no or negative feedback). (**c**) Feedback distribution curve with accessions carrying the alternative allele of the SNP at position 16,203,882 indicated with black lines and filled circles. (**d**) The genomic region (50 kb) flanking the SNP at position 16,203,882, including the two TNL genes *AT3G44630* (*MMF1*) and *AT3G44670*. (**e**) Venn diagram showing the intersection between the genes located closest to the top-ranking 100 SNPs in (**a**), and genes differentially expressed (false discovery rate < 0.05; empirical Bayes’ shrinkage, DESeq2) in the root and shoot transcriptome of Col-0 grown on the BXplus and BXminus microbiome (data from ref (*30*)).

### MMF1 mediates differential microbiome feedbacks

To validate the function of *MMF1* in microbiome growth feedbacks, we focussed on the reference accession Col-0 in which the positive feedback was first observed (**Fig. 1**). We phenotyped two mutant T-DNA insertion lines in Col-0 background for their response to the BX_minus_ and BX_plus_ microbiomes. At the transcriptional level, the *mmf1-1* and *mmf1-2* lines are knockdown and knockout mutants, respectively (**Extended Data Fig. 4**). Feedback assays were terminated at 35 DAG, when differential growth feedbacks were consistently visible in previous experiments with Col-0 (**Fig. 1**). Both mutants resembled the wild type in overall appearance - aside from slightly reduced size - and displayed no disease symptoms (**Fig. 4a**, **Dataset 4** and **Suppl. File 4**). Consistent with the initial experiments (**Fig. 1**, (*30*)) and the GWAS experiment (**Fig. 2c**), wild-type plants expressed positive growth feedbacks in response to the BX_plus_ microbiome, with higher fresh weight and larger rosette area compared to those growing on the BX_minus_ microbiome (**Fig. 4b**). In contrast, both *mmf1* mutants lost the positive growth feedback to the BX_plus_ microbiome, reflected in fresh weight and rosette area. This phenotype was reproducible in two additional experiments using independently conditioned soil batches (**Extended Data Fig. 5a, b**) and supported by a linear mixed-effects model (LMM) and estimated marginal means (EMM) across all three experiments (**Extended Data Fig. 5c**). The line-specific estimates confirmed that wild-type plants significantly responded to the BX_plus_ microbiome with enhanced fresh weight (LMM: *p* < 0.001; EMM: *β =* 0.036, *p* < 0.001) and rosette area (LMM: *p* = 0.023; EMM: *β =* 1.6 × 10^5^, *p* = 0.024), while the effect of the BX_plus_ response was attenuated in *mmf1-1* (fresh weight: LMM: *p* = 0.055; EMM: *β =* 0.006, *p* = 0.583; rosette area: LMM: *p* = 0.25; EMM: 4.1 × 10^4^, *p* = 0.587) and lost in *mmf1-2* (fresh weight: LMM: *p* = 0.033; EMM: 0.004, *p* = 0.698; rosette area: LMM: *p* = 0.089; EMM: 9.6 × 10^3^, *p* = 0.891). Complementary to the mutant analysis, we performed genetic complementation of the *mmf1-2* mutant using a functional *MMF1* allele driven by the nopaline synthase promoter. In these lines, the growth feedback was partially restored (**Fig. S2**, **Suppl. Results**), confirming that the loss of *MMF1* is responsible for the absence of the differential growth feedback in *mmf1* mutants. The genetic validation with mutant and complementation lines allowed to conclude that a functional MMF1 is required for the differential growth feedback of Col-0 in response to the two soil microbiomes.

**Figure 4.**
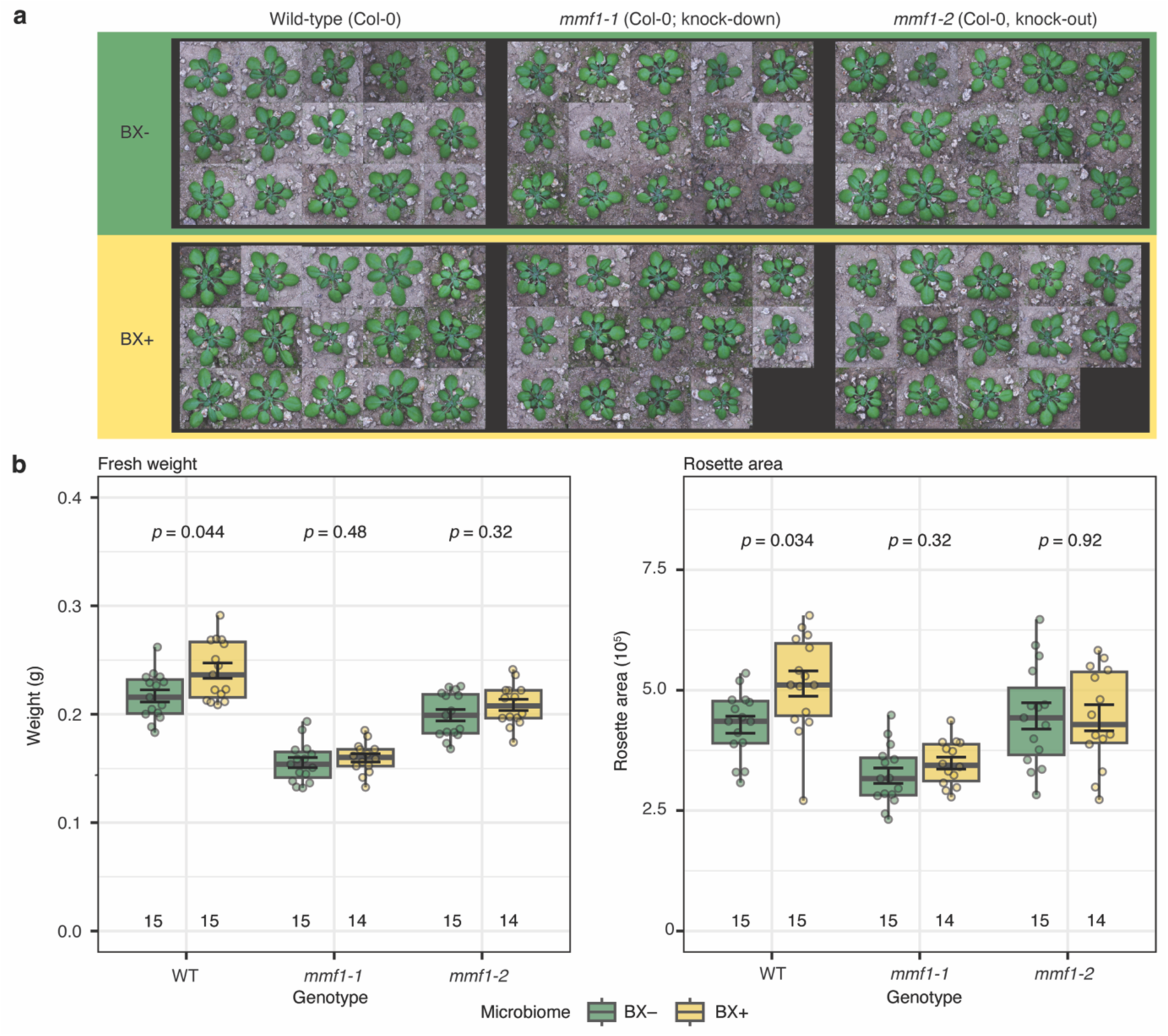
MMF1 is involved in growth feedback responses to the BXplus microbiome. (**a**) Representative rosettes of 6-week-old Col-0 (WT) and two independent *mmf1* T-DNA insertion mutants (*AT3G44630*) grown on BXminus and BXplus soil microbiomes (experiment IV, **Table S1**). Scale bar, 2 cm. (**b**) Fresh weight (left) and rosette area (right; in pixels) after 6 weeks of growth for the genotypes in (**a**). The findings were reproduced in two additional experiments (experiments III and V) on two separately conditioned soil batches. Differences were analysed with a two-way ANOVA (*line* × *microbiome*) followed by Benjamini–Hochberg FDR correction; adjusted *p*-values are reported. Sample sizes (n) are given below each boxplot. Points represent individual plants.

The plant immune system relies on two TNL signalling pathways: Enhanced Disease Susceptibility 1 (EDS1) together with Phytoalexin Deficient 4 (PAD4) lead to resistance, and EDS1 jointly with Senescence-Associated Gene 101 (SAG101) induce hypersensitive responses and cell death (*41*). To test the involvement of the two TNL signalling pathways in microbiome feedbacks, we performed feedback assays with *eds1*, *pad4* and *sag101* mutant plants (*42*, *43*). These mutants showed no altered growth phenotypes or obvious disease symptoms in response to BX_minus_ or BX_plus_ soil microbiomes (**Extended Data Fig. 6a**). Similar to wild type, the three immune signalling mutants expressed positive growth feedbacks in response to the BX_plus_ soil microbiome (**Extended Data Fig. 6b**). Hence, single-gene knockouts in EDS1, PAD4 and SAG101 do not disrupt immune signalling that leads to growth feedbacks, or alternatively, suggest that this process may be independent of these two classical pathways.

### MMF1 affects root bacterial community assembly sourced from the BXplus soil microbiome

In earlier work, we found differential microbiomes on roots of plants with differential growth feedbacks (*14*, *30*). Because *MMF1* encodes a defence-related protein, we profiled the root bacterial, fungal and oomycete communities recruited by wild-type and *mmf1* mutant plants when grown in BX_minus_ and BX_plus_ soils. The detailed microbiome analysis is documented in **Dataset 5** and **Suppl. File 5**. We utilised hamPCR (*44*), which determines microbial load relative to host tissue, for microbiome analysis and noticed in factorial ANOVA of microbial loads that the interaction term between the *line* (WT and *mmf1-2*) and *microbiome* (BX_minus_ and BX_plus_) factors was close to significance for bacteria, but not for fungi or oomycetes (**Extended Data Fig. 7a**). Bacterial load was significantly higher in the roots of wild-type plants compared to *mmf1* mutants on the BX_plus_ microbiome. While alpha diversity (measured with Shannon index) did not differ between the factors *line* (WT and *mmf1-*2) and *microbiome* (BX_minus_ and BX_plus_; **Extended Data Fig. 7b**), they impacted the compositions of the root microbial communities (beta diversity tested with Permutational Multivariate Analysis of Variance (PERMANOVA)). Source *microbiome* accounted for little variation in bacterial (*R^2^ =* 4.6%, *p* = 0.059), fungal (*R^2^ =* 3.0%, *p* = 0.25) and oomycete communities (*R^2^ =* 4.7%, *p* < 0.05; **Fig. 5a**). For *line* however, we found significant differences in bacterial – but not fungal or oomycete – communities between wild type and *mmf1* mutants (*R^2^* = 5.9%, *p* < 0.05) and also in their interaction of the factors *line* with *microbiome* (*R^2^* = 4.5%, *p* < 0.05). The interaction effect indicated that the root microbiome of the mutant differed from wild type in only one soil type. Pairwise PERMANOVA confirmed this with more and significant variation between wild type and mutants in the BX_plus_ (*R^2^* = 18.3%, *p* < 0.01) compared to the BX_minus_ (*R^2^* = 5.1%, *p* = 0.548) soil microbiome. In contrast, fungal and oomycete communities showed no differences between *line* and *microbiome* (**Extended Data Fig. 7b**). Constrained Analysis of Principal Coordinates (CAP) confirmed these findings: root bacterial communities clearly separated between wild type and mutant on the BX_plus_ but not the BX_minus_ soil microbiome (**Fig. 5b**), while fungal and oomycete communities showed minimal, non-significant separation (**Extended Data Fig. 7c**). Taken together, the *mmf1* mutant assembled a distinct bacterial root microbiome relative to wild-type plants, but only on the BX_plus_ source microbiome.

**Figure 5.**
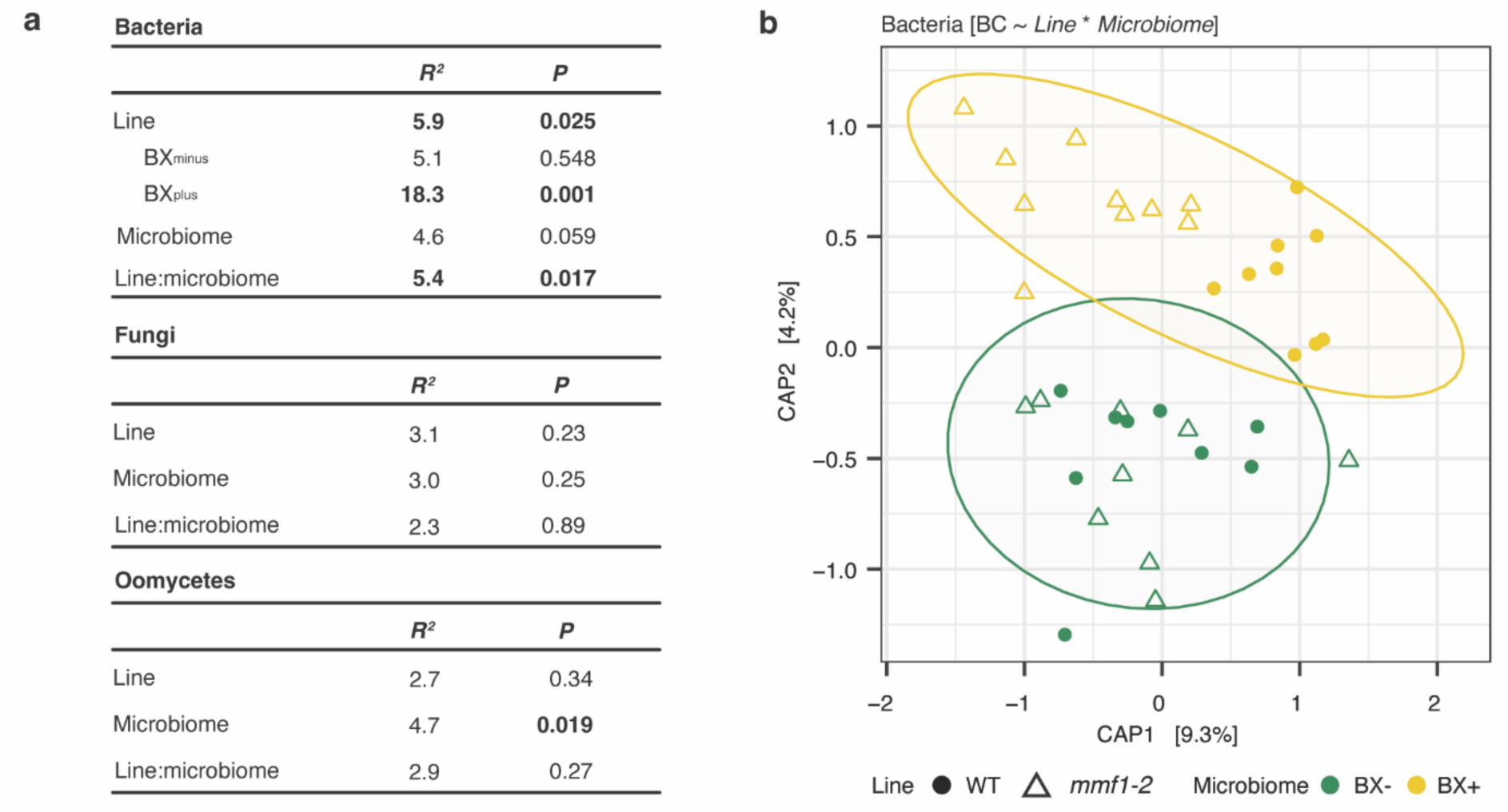
Root microbiome profiling of wild-type and *mmf1-2* mutants grown on BXplus or BXminus soil. (**a**) The output of PERMANOVA on Bray-Curtis dissimilarities of bacterial, fungal and oomycete root microbiome of Arabidopsis wild-type (WT; Col-0) and *mmf1-2* mutants grown on the BXminus or BXplus soils (sampled from experiment IV, **Table S1**). The *R^2^* and *p-*values indicate the *line* and *microbiome* (conditioning) effects and their interaction. Pairwise PERMANOVA for bacterial communities compares WT and *mmf1-2* mutants on each soil condition; significant effects (*p* < 0.05) are indicated in bold. (**b**) Constrained Analysis of Principal Coordinates (CAP) of Bray-Curtis dissimilarities [BC] for WT (filled circles) and *mmf1-2* mutant samples (open triangles) grown on the BXminus (green) or BXplus (yellow) soils confirming the effects of the soil conditioning and lines observed in the PERMANOVA analysis. Ellipses represent 95% confidence intervals for plants grown on BXminus (green) or BXplus (yellow) soils. Axes show the percentage of variance explained by the first two principal coordinates.

To identify the taxa that differed in abundance between roots of wild type and *mmf1* mutants across the two soil microbiomes, we used DESeq2 (*45*) for modelling the same factors *line*, *microbiome*, and their interaction. We retained only amplicon sequence variants (ASVs) with a false discovery rate (FDR) < 0.05 for any model term. A total of 26 ASVs were differentially abundant: 18 by *microbiome*, 12 by *line*, and 20 by their interaction (**Extended Data Fig. 8**). ASVs assigned to *Pelomonas*, *Massilia*, and *Asticcacaulis* were more abundant taxa in roots from plants grown in BX_plus_ soils. *Lechevalieria*, *Variovorax*, and *Duganella* differed between genotypes and showed significant interaction effects. *Pseudomonas*, *Lechevaleria* and *Aminobacter* were the most dominant ASVs, differing only by the interaction term. We noticed that the latter all showed reduced abundance levels in the roots of *mmf1* mutants grown on the BX_plus_ microbiome compared to wild type.

### MMF1 is required for a defence-related transcriptional response

To obtain first insights into how MMF1 contributes to positive growth feedbacks, we analysed the root transcriptomes of wild-type and *mmf1-2* mutants in response to both soil microbiomes (detailed transcriptome analysis documented in **Dataset 6** and **Suppl. File 6**). Analogous to the microbiome analysis, we first examined the contributions of the factors *microbiome* and *line* on the transcriptome using PERMANOVA. *Microbiome* accounted for 29.0% (based on *R^2^*, *p* < 0.01), *line* for 33.7% (*p* < 0.01), and their interaction for 12.4% (*p* < 0.01) of the transcriptomic variation (**Fig. 6a**). Again, the significant interaction term revealed that mutant and wild type responded differently to the two soil microbiomes. Pairwise PERMANOVA again revealed greater variation in wild type (*R^2^* = 79.7%, *p* < 0.01) compared to mutant (*R^2^* = 10.8%, *p* = 0.349) plants in response to the two microbiomes. Principal Component (PC) analysis confirmed a clear separation of wild-type individuals between the two source microbiomes (PC1, 48% of variation) and between plant lines (PC2, 16% of variance; **Fig. 6b**). Overall, the strongest separation was between wild-type plants in response to the two microbiomes, whereas the *mmf1* mutant transcriptomes clustered together.

**Figure 6.**
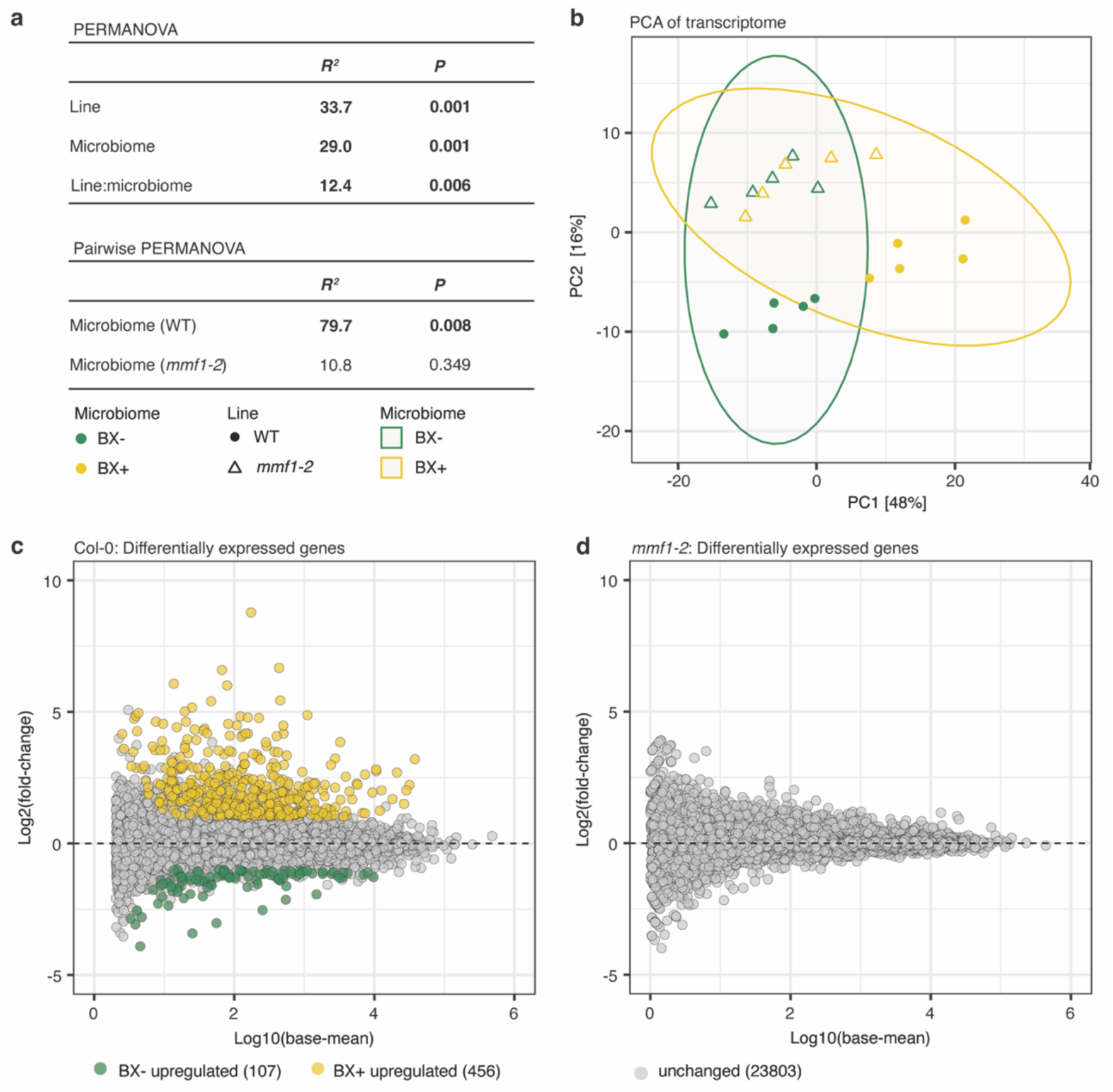
Plants lacking MMF1 fail to mount a transcriptional response to a BX microbiome. (**a**) The output of PERMANOVA on Euclidean distances of the root transcriptome of wild-type (WT) and *mmf1-2* mutant plants in response to the BXminus and BXplus microbiomes (sampled from experiment IV, **Table S1**). The *R^2^* and *p-*values summarize the effects of *line* (plant genotype) and *microbiome* (soil condition) along with their interaction. Pairwise PERMANOVA results compare the transcriptomes of WT and *mmf1-2* mutant plants when grown on the BXminus soil microbiome compared to the BXplus soil microbiome; significant effects are indicated in bold. (**b**) Principal component analysis (PCA) of the same root transcriptome confirms the *line* (WT, closed circles; *mmf1-2*, open triangles) and *microbiome* (BXminus, green; BXplus, yellow) effect of the PERMANOVA results. The axes display the percentage of variance explained by each principal component. (**c**) MA plots illustrating differentially expressed genes (DEGs) in WT between BXminus and BXplus (left), and the absence of DEGs in *mmf1-2* (right). Genes upregulated on the BXplus microbiome (yellow) or BXminus microbiome (green) are colour coded. Genes with an absolute log2 fold-change ≥ 1 and false discovery rate < 0.05 (empirical Bayes’ shrinkage; DESeq2) are considered significant.

Next, we analysed the transcriptomes at the gene level to determine the differentially expressed genes (DEGs; log_2_ fold-change ≥ 1; empirical Bayes shrinkage, *p* values adjusted for FDR < 0.05). Aligned with the microbiome analysis (**Fig. 5**), we found more DEGs between the lines in response to the BX_plus_ (347 total; 230 up and 117 down in wild type) compared to the BX_minus_ soil microbiome (128 total, 68 up and 60 down; **Extended Data Fig. 9a, b**). Wild-type plants showed 563 DEGs in response to the two microbiomes (107 down and 456 up; **Fig. 6c**). Surprisingly, not a single *microbiome*-specific DEG was found in mutant plants (**Fig. 6d**). Taken together, a functional MMF1 leads to a strong transcriptional reprogramming, mostly in response to the BX_plus_ soil microbiome.

Finally, we performed co-expression and gene ontology (GO)-term enrichment analysis (underlying genes documented in **Dataset 6**) to characterise the transcriptional changes of wild-type plants in response to the two microbiomes (**Fig. 6c**). Hierarchical clustering identified clusters of co-expressed genes: four up- and three downregulated (**Fig. 7**, **Extended Data Fig. 9c**). Upregulated processes included hypoxia (clusters #2 and #3) and immune responses (clusters #2 to #5). Processes linked to hypoxia included the GO terms response to hypoxia and cellular response to hypoxia (**Dataset 6**). Upregulated immune processes included GO terms such as response to ethylene and systemic acquired resistance (cluster #2), defence response to insects, and response to bacterial molecules (cluster #3), indole metabolism and response to salicylic acid (SA; cluster #4), and hypersensitive response, programmed cell death, and interaction with symbionts (cluster #5; see **Dataset 6** for the genes in each cluster). Of note, several of these genes are typically involved in TNL-triggered immunity (*41*, *46*). Downregulated processes included fatty acid biosynthesis, secondary metabolite biosynthesis and malate transport (clusters #6 to #8; **Dataset 6**). In conclusion, wild-type plants, which achieved better growth in response to the BX_plus_ soil microbiome, exhibited a strong transcriptional reprogramming, most noticeably an upregulation of defence-related genes.

**Figure 7.**
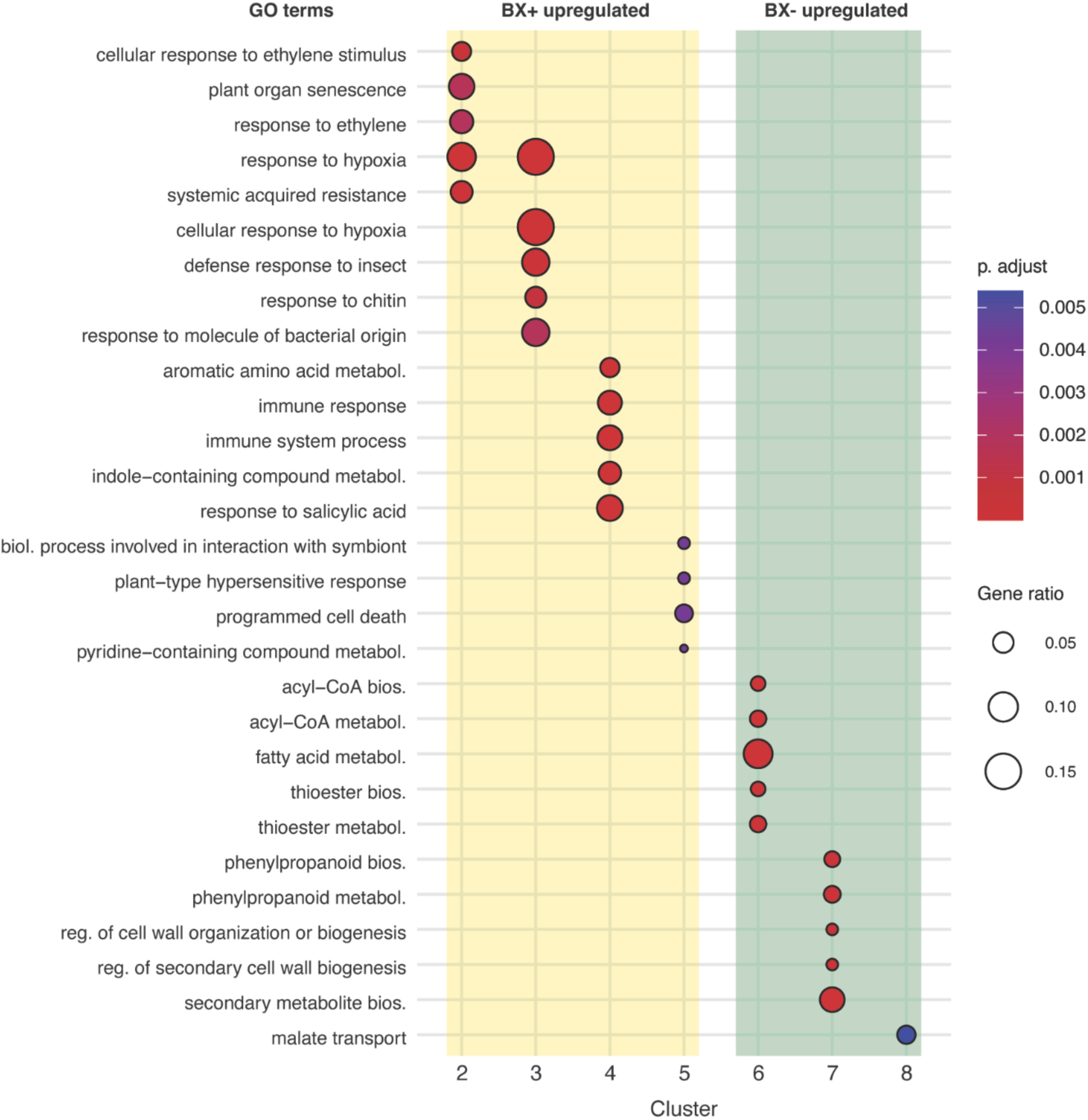
Gene ontology term enrichment reveals plant immune responses in response to the BXplus microbiome. Hierarchical clustering and gene ontology (GO) term enrichment analysis performed on differentially expressed genes in wild-type roots grown on the BXplus relative to BXminus soil microbiomes. The top five most significantly enriched GO terms (Benjamini–Hochberg, *p* < 0.01) for each cluster are shown. Point size indicates the ratio of genes in a cluster mapping to the respective GO term, and point colour reflects the adjusted *p*-value (FDR corrected using Benjamini-Hochberg). Shading indicates clusters upregulated in plants grown on the BXplus microbiome (yellow) or upregulated in plants grown on BXminus microbiome (green). No significantly enriched GO terms were found for clusters 1 and 9. Bios., biosynthesis; metabol., metabolism; boil., biological; reg., regulation.

## Discussion

While genotype-dependent microbiome feedbacks have been observed in many plant systems (*14*, *28*, *29*, *33*), the mechanisms underlying plant responses to complex microbiomes remain largely unresolved. Using our model system with Arabidopsis, where the reference accession Col-0 expresses differential growth responses on two different soil microbiomes (*30*), we set out to uncover the genetic basis of these microbiome feedbacks. Here, we used natural variation in growth feedbacks to BX_plus_ and BX_minus_ soil microbiomes among natural Arabidopsis accessions (**Fig. 2c**) as a tool to uncover candidate loci through which plants respond to different microbiomes in natural soils. We focussed on the gene *AT3G44630*, which was in the proximity of the top-ranking SNP in the GWAS (**Fig. 3**), named this TNL-receptor-encoding gene *MMF1*, and characterised it in the reference accession Col-0. Structural modelling revealed that the predicted MMF1 tetramer shares high similarity with the tetrameric resistosome structure of the well-characterized TNL RPP1 (*40*), including conserved TNL domains and tetrameric assembly (**Fig. S1**). These predictions suggested that MMF1 may function as a TNL perceiving microbial signals. In Col-0, *mmf1* mutants lacked three hallmarks of wild-type plants in response to the BX_plus_ soil microbiome: they lost the positive growth feedback (**Fig. 4**), had an altered root bacterial community and bacterial load (**Fig. 5**), and failed to trigger the defence-related transcriptional response (**Fig. 6**). Given the molecular function of TNLs – to recognise microbially-derived signals intracellularly and trigger an immune response (*20*) – our results support the following mechanistic model (**Fig. 8**): When grown on the BX_plus_ soil microbiome, wild-type plants show enhanced shoot growth resulting from a defence-related transcriptional response (**Fig. 7**). This MMF1-dependent transcriptional response, together with the typical intracellular localisation of TNLs, suggest that a bacterial (see below) signal of the BX_plus_ microbiome is perceived inside the root cells. This model is consistent with the reduced shoot growth of wild-type plants on the BX_minus_ microbiome (in the absence of the signal) as well as *mmf1* mutants on the BX_plus_ microbiome (lack of the receptor), and is further supported by the lack of a defence-related transcriptome in *mmf1* mutants.

**Figure 8.**
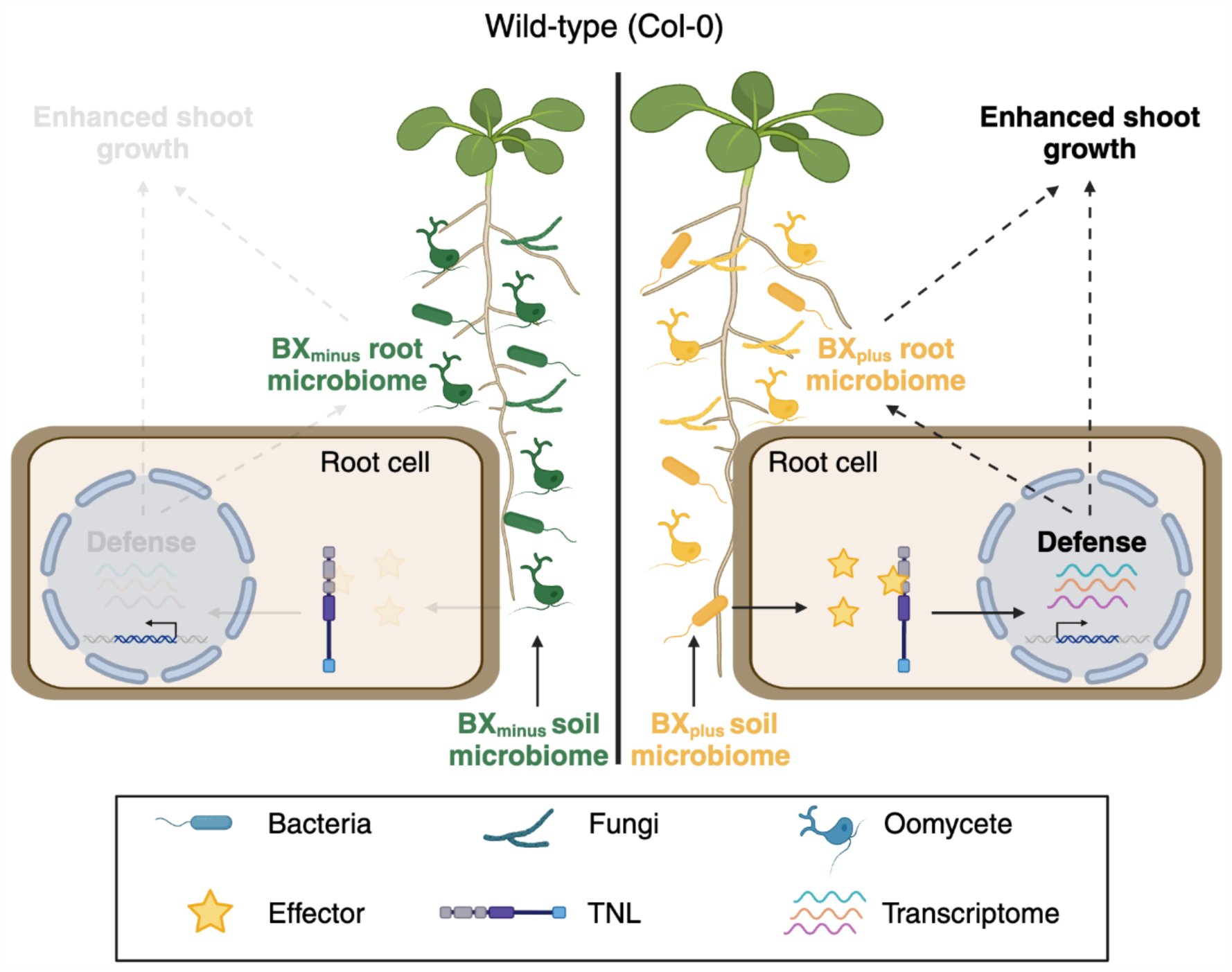
Proposed model of microbiome feedbacks through MMF1 in Col-0 plants. Plants with a functional MMF1 receptor perceive a putative effector molecule that is present in the BXplus soil microbiome, mounting a defence response that coincides with an altered root microbiome and promoted plant growth. The absence of this effector molecule in the BXminus microbiome results in the lack of defence responses in plants grown on this soil microbiome. Col-0 plants lacking the MMF1 receptor fail to perceive the effector molecule from the BXplus microbiome and do not mount defence responses; they have a different microbiome, and no enhanced shoot growth is observed. Figure created with BioRender^®^.

The plant responses, including a defence-related transcriptome and enhanced shoot growth, coincided with an altered root microbiome (**Fig. 5**). While, overall, root microbiomes were similar between soil sources, the bacterial load and communities of wild-type and *mmf1* mutant plants differed significantly on the BX_plus_ but not on the BX_minus_ microbiome (**Fig. 5**). These data do not allow to establish a causal link between the altered root microbiome and the differential shoot growth of the genotypes. However, the observation suggests that a *bacterial* signal in the BX_plus_ microbiome, which presumably triggers the defence-related transcriptome via the TNL MMF1, may be responsible. Isolation of the bacterial species, possibly guided by the analysis of differentially abundant ASVs (**Extended Data Fig. 8**) and the actual signal (i.e. a putative effector) are obvious future research targets.

As part of the immune system, TNLs act as gatekeepers against opportunistic pathogens, maintaining host health and microbiome homeostasis (*20*). Earlier work showed that Arabidopsis with impaired ETI undergo microbiome dysbiosis in their leaves, and ultimately these plants have reduced fitness (*24*, *47*). Analogously, in roots, we report microbiome dysbiosis in the mutant *mmf1* and demonstrate that MMF1 is required for growth promotion in wild-type plants. Both examples imply that TNLs integrate microbiome signals inside host cells for maintaining microbiome homeostasis and host health. While TNLs and their signalling pathways have been predominantly studied in single-pathogen and gene-for-gene interactions, this model faces a multiplicate challenge at the microbiome level, with many more signals being present in a diverse microbiome (*16*). Thus, plant immune responses rely on more complex signal integration when interacting with microbial communities compared to single isolates (*16*, *47*). Moreover, there is evidence that signal integration may further depend on TNL receptor dosage: recently, high TNL copy numbers within the *RPP-1*-like gene cluster among Arabidopsis accessions were linked with reduced bacterial diversity in their phyllosphere (*48*). Of note, the closest homolog of *MMF1* is *RPP1* (*49*), and additional TNL encoding genes are in the proximity of the SNP most strongly associated with the variation in growth feedback among the tested accessions (**Fig. 3d**). Here, we have shown that lack of MMF1 is sufficient for the loss of the previously established growth feedback in Col-0, but future work is needed to test whether additional members of the TNL family or their dosage (*48*) also contribute to feedback in other accessions.

We set out to identify a first genetic component through which plants respond to different microbiomes in natural soils. The substantial variation in growth feedbacks among Arabidopsis accessions (**Fig. 2c**) implies that there are additional components to discover. *MMF1* presents a first genetic component for such feedbacks, but for now it only explains the enhanced growth and increased defence of the reference accession Col-0 responding to the BX_plus_ microbiome. MMF1 activity alone cannot explain the massive variation in growth feedbacks among the other accessions. For instance, Hadd-3, which shares the reference allele of the top SNP with Col-0, grew more poorly on the BX_plus_ microbiome, while IP-Hue-3 harboured the alternative SNP allele and grew much better than Col-0. Solving this puzzle requires further analysis of the complex locus around the top SNP, and investigating MMF1 variation of the extreme accessions such as Hadd-3 and IP-Hue-3. Together with the other GWAS candidate loci (**Fig. 3a**), additional genetic components likely contribute to explaining the massive variation in growth feedbacks among the accessions. It is also possible that the genetic architecture of the identified locus shows a more complex, for example structural, variation pattern across the study population.

In conclusion, the identified TNL receptor provides a relatively straightforward explanation for the enhanced shoot growth of Col-0 on the BX_plus_ soil microbiome: MMF1 perceives a signal of the BX_plus_ microbiome, triggers defences, and alters the root microbiome, which collectively coincide with better shoot growth mediated by yet unknown mechanisms. This work contributes to understanding host genetic pathways by which plants perceive complex soil microbiomes and mediate microbiome feedbacks. Such knowledge may pave the way towards crops that leverage beneficial microbiome feedbacks, ultimately improving yield in a more sustainable manner.

## Methods

### Plant material

The maize (*Zea mays* L.) genotypes B73 and *bx1* (B73) transposon insertion mutant (*50*) were used for soil conditioning (see below). The 410 *Arabidopsis thaliana* (hereafter: Arabidopsis) accessions used in the GWAS (see below) were obtained from the 1001 genomes project (The 1001 Genomes Consortium, 2016). For a full list of the accessions used, see **Dataset 1**. Arabidopsis T-DNA insertion mutants (AT3G44630) for *mmf1-1* (SALK_144159C) and *mmf1-2* (SALK_002318C) are in the Col-0 background and were obtained from the Nottingham Arabidopsis Stock Centre (NASC). Mutants were confirmed for homozygosity and characterised by qPCR (**Extended Data Fig. 4**; primers in **Table S3**). The mutant lines *eds1-12* (*42*), *pad4-1* (*51*), and *sag101-3* (*52*) in Col-0 background were kindly provided by Prof. Jane Parker.

### Generation of BX_minus_ and BX_plus_ soil microbiomes

We used soil from an agricultural field site of Agroscope in Changins, Switzerland (46°24’00.0“N6°14’22.8”E; three adjacent plots, Parcel 29, Parcel 30, or Parcel 31) and we generate the distinct soil microbiomes either in the field or in the greenhouse using pots filled with that soil. Starting from replicates of the same batch of unconditioned field soil, the two distinct microbiomes are generated by growing of either wild-type maize (B73), which secrete benzoxazinoids (BXs) to the surrounding soil, or BX-deficient mutant maize (*bx1* (B73)). After three months of growth, the plants and roots are removed and the two maize lines “leave behind” BX-conditioned (BX_plus_) and control (BX_minus_) microbiomes in the soil. We then harvest either replicate soil cores (20 × 20 × 20 cm, conditioning in the field) or soil from replicate pots (3 L, 13.1 cm deep and 20.2 cm diameter; conditioning in the greenhouse), sieve the soil through a 1 cm^2^ sieve and stored it at 4°C until the subsequent feedback experiments. We described in detail the approach of generating BX_minus_ and BX_plus_ soil microbiomes (we often referred to this as ‘soil conditioning’) including the physicochemical characteristics of the soil (*14*, *28*, *38*), management practices and cropping history of the field site (*14*, *26*) and the conditions for greenhouse conditioning (*14*, *28*, *30*). The batches of conditioned soils for this study, and the experiments linked to soil batches are listed in **Table S1** and **S2**.

### Microbiome feedback experiments

We list the setups and purposes of different experiments (Exp. I to Exp. VII) with Arabidopsis grown on BX_minus_ and BX_plus_ soil microbiomes in **Table S2**. For feedback experiments, the replicate soil samples containing the BX_plus_ and BX_minus_ microbiomes were individually amended with sand, by manually mixing in 20% (v/v) autoclaved quartz sand (*14*). Pots were equally filled by weight and randomized across trays. Below we detail the specific setups and conditions for the high-throughput phenotyping of the different Arabidopsis accessions (Exp. I and II) and the characterisation of wild-type and mutant lines in Col-0 background (Exp. III-VII).

#### High-throughput phenotyping of Arabidopsis accessions

We performed two microbiome feedback experiments; each experiment was done with 210 accessions (10 accessions repeated in both runs) resulting in a total of 410 tested Arabidopsis genotypes (**Dataset 1**). In each experiment, every accession was seeded on both BX_minus_ and BX_plus_ soils with 3 replicates (2 experiments of 210 accessions x 2 soils x 3 replicates = 1260 pots). Plant cultivation and phenotyping was previously described and the images of the first experiment were utilized to develop the phenotyping software ARADEEPOPSIS (without comparing BX_minus_ and BX_plus_ soils) (*35*). In brief, plants were grown at the Gregor Mendel Institute of Molecular Plant Biology, Vienna, Austria, in a custom-built growth chamber (YORK Refrigeration), equipped with an environmental control unit (Johnson Controls) and a climate control unit (Incotec) under long-day conditions (16h light [21°C], 8h dark [16°C], 140 µE/m^2^s) with 60 % relative humidity and watered twice per week by flooding. Plants were imaged twice daily, and plant morphometric phenotypes extracted using the phenotyping software ARADEEPOPSIS (*35*). The phenotypes were converted into “days after germination” phenotypes by taking the average of the two measurements that happened on the same day. **Dataset 1** contains the source data and **Suppl. File 1** documents the analysis.

#### Feedback experiments with Arabidopsis mutants

For microbiome feedback experiments performed on Arabidopsis Col-0 mutant lines, we used greenhouse conditioned soils prepared as described before. Seeds were stratified for 3 days in the dark at 4 °C and sowed on pots (5.5 × 5.5 × 5 cm) filled by weight. Plants were grown in a plant growth chamber (Percival, CLF Plant Climatics, Wertingen, Germany) under short days with 10h light (100 μmol m^−2^ s^−1^) (21 °C) and 14h darkness (18 °C) at 60% relative humidity. Plants were fertilized at day 14 and 21 after germination using 10 mL of 1:3 times diluted ½ strength Hoagland solution (*53*), watered 3 times per week by weight, and randomized weekly. Analysis was blinded by adding barcodes to the pots for phenotyping which was used to trace back sample names after analysis. Plants were photographed weekly and phenotyped using the ARADEEPOPSIS pipeline to obtain the rosette area (plant region area) (*35*). Plants were grown for 6 weeks before the rosettes and roots were harvested for downstream analysis. **Dataset 4** contains the source data and **Suppl. File 4** documents the analysis.

### Genome-wide association study

Genome-wide association studies (GWAS) search for associations between single nucleotide polymorphisms (SNPs) in a genome and a phenotype (*54*). As phenotype we took the differential growth feedback of the accessions on the two soil microbiomes, i.e. the mean growth on BX_minus_ was subtracted from the mean growth on BX_plus_ soil microbiomes [BX_plus_ – BX_minus_]. We focussed on the first 20 days after germination (DAG) to avoid confounding effects of phenotypes like early flowering and overlapping leaves. GWA was performed using an in-house Nextflow pipeline (https://gitlab.lrz.de/beckerlab/gwas-nf) that wraps limix (*39*) and applies the same multi-trait model used previously in (*35*). Only SNPs with a minor allele frequency of at least 1% were included. The model estimates each SNP’s “specific” effect on the trait “differential growth feedback”. In this study, we used plant_region_area as a trait (**Dataset 1**), and the specific effect reflects the difference in that measurement between the soil conditions. The analysis was conducted independently for each day because we noted temporal variation in daily growth feedbacks, and the resulting tables were combined without re-adjusting the corrected p-values. SNPs were retained and visualized if they had a −log_10_(adjusted p-value) of at least 5 on at least three consecutive days and a minor allele count of at least 5. From these plots, we identified SNPs with a robust multi-day signal and analysed them further using gwaR (https://github.com/Gregor-Mendel-Institute/gwaR). This allowed us to investigate the genomic neighbourhood of those SNPs and pinpoint genes likely involved in the microbiome feedback phenotype. The GWAS data are in **Dataset 3** and their analysis is documented in the **Suppl. File 3**. For computational analyses, we used the BioHPC-Genomics compute cluster at the Leibniz Rechenzentrum and the CLIP cluster at Vienna BioCenter (VBC).

### GWAS candidate gene selection

To identify candidate genes associated with BX-growth feedbacks, we performed the genome wide association study (GWAS) with the 410 genome-sequenced accessions using their phenotypic diversity in growth feedbacks (**Fig. 2c**). To account for population structure within the *Arabidopsis* accessions, a linear mixed model was employed (*39*). We used the daily growth feedback data, which includes the temporal dynamics (**Extended Data Fig. 2b**), and calculated associations between single nucleotide polymorphisms (SNPs) and phenotypes for each of the 20 DAGs (**Dataset 3**). In essence, we performed the GWAS using each day as a covariable in the analysis.

To impose stringency in SNP identification, we performed FDR correction and Bonferroni correction of the *p*-values of SNPs that associated with the growth feedbacks (**Fig. 3a**, **Dataset 3**). Next, we filtered the SNPs for those with a minor allele frequency of higher than 2.5 % to also include rare SNPs associated with local and environmental adaptation (*55*), and those SNPs with significance (Bonferroni correction) for at least three consecutive days. To isolate candidate genes, we used the “get_nearest_genes” function from the gwaR package in R (https://github.com/Gregor-Mendel-Institute/gwaR) to identify genes located near each candidate SNP. Finally, to select candidate genes for functional validation, we screened the genes identified in the GWAS for overlap with differentially expressed genes in the roots and shoots of Arabidopsis plants grown on a BX_minus_ and BX_plus_ microbiome (*30*).

### Validation of single T-DNA insertion in mmf1-2

We validated that *mmf1-2* carries a single T-DNA insertion by long-read whole genome sequencing and reference-guided assembly. High molecular weight DNA was extracted from leaves using the Machery-Nagel NucleoBond HMW DNA kit according to manufacturer’s instructions. Long-read sequencing was carried out on an Oxford Nanopore Technologies promethion 24 instrument in a multiplexed run on a R10.4 flow cell. After demultiplexing with dorado (https://github.com/nanoporetech/dorado) adapters were trimmed using porechop, and the genome was assembled using flye (*56*), polished with medaka (https://github.com/nanoporetech/medaka) and scaffolded against Col-CEN (*57*) using ragtag (*58*) and annotations (araport11) from Col-CEN were lifted over using liftoff (*59*). The T-DNA insertion in AT3G44630 was identified by aligning to the reference genome. The T-DNA sequence was used as a query for BLAST against the *mmf1-2* assembly, returning only the insertion in AT3G44630 (**Extended Data Fig. 4d**).

### Generation of MMF1 complementation lines

To generate *MMF1* complementation lines in the *mmf1-2* mutant background, we used a binary vector containing the full-length *Arabidopsis thaliana MMF1* gene (*AT3G44630*, transcript ID: NM_180324.2) under the control of the nopaline synthase (NOS) promoter. The expression construct was commercially synthesized and cloned by VectorBuilder (vector name: pPBV[Exp]-Hygro-Nos>ath_AT3G44630[NM_180324.2]) and includes a hygromycin resistance cassette for plant selection. The construct was introduced into *Agrobacterium tumefaciens* strain AGL-1 by electroporation. Arabidopsis *mmf1-2* homozygous mutants were transformed using the floral inoculation method (*60*). Transformed seeds were selected on Murashige and Skoog (MS) medium supplemented with hygromycin (20–30 mg/L). Resistant T1 seedlings were transferred to soil, and independent transgenic lines were screened for *MMF1* expression (**Fig. S2c**) and T3 lines used in downstream phenotypic analyses.

### MBOA allelopathic growth assays

To test for the allelopathic effect of MBOA on plant growth, we grew different Arabidopsis accessions on ½ MS (Duchefa, Haarlem, the Netherlands) plates containing 1.5% phytoagar (Duchefa, Haarlem, the Netherlands) supplemented with 0.25 mM MBOA (Sigma Aldrich, St. Louis, USA) dissolved in DMSO or on control plates supplemented with the same concentration (500 μL per L) of DMSO (*61*). Seeds were stratified by incubating plates for 3 days at 4 °C. The plates were placed upright and grown in a controlled growth chamber (Percival, CLF Plant Climatics, Wertingen, Germany) in short days with 10 hours light (100 μmol m^−2^ s^−1^) (21 °C) and 14 hours darkness (18 °C) at 60 % relative humidity. After 10 days, the lateral root length was measured using the image software ImageJ (*62*).

### Root microbiome profiling

#### DNA extraction

For root microbiome analysis, 10 randomly selected *Arabidopsis* root systems were removed from the pots, and excess soil carefully removed by hand. The root system was rinsed with sterile ddH_2_O to remove additional soil. The entire root system was flash frozen in liquid nitrogen and stored at -80 °C until further processing. The roots were lyophilized for 24 h and grinded in 1.5 mL Eppendorf tubes using a Retsch mill (Retsch GmBH, Haan; Germany) and 2 mm glass beads at 30 Hz for 1 min. Approximately 20 mg of dry root material was used as the input for DNA extractions. DNA extractions were performed in 96-well plates using the QIAGEN® MagAttract® PowerSoil® DNA KF Kit (Qiagen, Germany) and the KingFisher™ Flex™ Purification System (ThermoFisher Scientific, Waltham, MA) following the manufacturers guidelines. For all samples, the DNA concentration was determined using the AccuClear® Ultra High Sensitivity dsDNA Quantitation Kit (Biotum, Fremont, United States). For library preparation, DNA concentration was adjusted to 2 ng/µL with a Myra Liquid Handler (Bio Molecular Systems, Upper Coomera; Australia), and 10 ng used for library preparation.

#### Library preparation

Library preparation for bacteria, fungi and oomycetes was performed using an adapted two-step barcoded host-associated microbiome hamPCR protocol (*44*). Primers used are documented in **Table S3**. For bacteria, the first PCR was performed using a combination of the CS1-*fs*-799-F and CS2-*fs*-1193R frameshift (fs) (pooled in 5 equimolar concentrations) in combination with the host specific *GIGANTEA* CS1-At-GI-F and CS2-At-GI-R primers in a single well. We followed earlier work PCR settings for annealing temperature (*63*) and cycling conditions (*44*). Briefly, the PCR settings were 94 °C for 2 min, and 10 cycles of 94 °C for 30 s, 55 °C for 50 s, 72 °C for 30 s and finally 72 °C for 5 min. For fungi, the first PCR was performed with the internal transcribed spacer (ITS) region primers ITS1fF and ITS2R with standard temperature (*64*) and cycle (*44*) settings. The PCR settings were 94 °C for 3 min, and 10 cycles of 94 °C for 45 s, 50 °C for 60 s, 72 °C for 90 s, and finally 72 °C for 10 min. For oomycetes, we adapted the ITS1-O_F1_G-46636 and 5.8s-O_R1_G-46637 primers (*44*) with the CS1 and CS2 adapter sequences and utilised the same PCR settings as for the fungi. PCR amplicons were purified using the ChargeSwitch® PCR Clean-Up Kit (Invitrogen, Waltham, MA, USA) on a KingFisher™ Flex™ Purification System (ThermoFisher Scientific, Waltham, MA, USA). The second PCR added unique barcodes to 5 µL of purified DNA using the Fluidigm Access Array barcoding primers (Standard BioTools, California, USA). The cycling conditions were 94 °C for 2 min, 25 cycles of 94 °C for 30 s, 60 °C for 30 s, 72°C for 60 s, and finally extension 72°C for 10 min. Samples were again purified as before and quantified and pooled by adding 80 ng of each sample using a Myra Liquid Handler to create sub-libraries for bacteria, fungi and oomycetes. The pooled bacterial sub-library was subsequently separated on an 1.5 % agarose gel, the bacterial and GIGANTEA fragments excised, and gel purified using the Wizard® SV Gel and PCR Clean-Up System (Promega, Madison, Wisconsin, USA). The final bacterial sub-library was created by scaling the purified bacterial and GIGANTEA amplicons in a 24:1 ratio (bacteria:GIGANTEA) to reduce the amount of plant reads during sequencing. Five random samples were duplicated during library preparation and left unscaled to bioinformatically rescale the scaled samples (see ref (*44*) for details). The final sequencing library was created by pooling the sub-libraries in a DNA ratio of 87.5% (963 ng) scaled bacteria, 5.5 % (60 ng) unscaled bacterial samples and 3.5 % (39 ng) each for fungal and oomycetes. The library was paired end sequenced on an Illumina MiSeq instrument (v3 chemistry, 300 bp paired end) at the NGS platform (https://www.ngs.unibe.ch) of the University of Bern.

#### Bioinformatics

MiSeq reads were quality checked with FastQC v0.11.8 (Babraham Institute, Cambridge, United Kingdom). Barcodes were removed previously by the NGS platform and written to the sequence headers. With this information the data was demultiplexed and primers were removed using cutadapt v3.4 (*65*). Following the methods from (*29*), quality filtering, read merging and amplicon sequence variant (ASV) inferring was performed with the dada2 package v1.26.0 (*66*) in R v4.2.0 (R Core Team 2022). Taxonomic assignment were done using naïve Bayesian classifier with a DADA2 formatted version from the SILVA v. 132 database (*67*) for bacteria and the FASTA general release from UNITE v8.3 for fungi. To improve taxonomy assignment, the taxonomy was assigned again with the IDTAXA classifier using the SILVA r138 and UNITE 8.2 databases from DECIPHER v2.26.0 (*68*) and replaced wherever IDTAXA assigned more taxonomic levels than the naïve Bayesian classifier. Bacterial ASVs assigned as Eukarya, Archaea, Cyanobacteria or Mitochondria and fungal ASVs belonging to Protista, Plantae, Protozoa or Animalia were discarded. Calculations were performed at the sciCORE (http://scicore.unibas.ch/) scientific computing centre of the University of Basel.

#### Statistical analysis

Statistical analyses were performed with the vegan package v.2.6.4 (*69*) in R v4.2.0. Following the strategies described before (*70*), we normalised the data for bacteria, fungi and oomycetes by rarefication. Rarefaction thresholds per sample were defined as 65’000 for bacteria, 400 for fungi and 5’700 for oomycetes (*70*). Outlier detection was performed with CLOUD using Bray-Curtis (BC) distances from the rarefied data for each condition and each treatment individually (*71*). The nearest number of neighbours were set to 60% of the sample size and we chose an empirical outlier percentile of 0.1. As described before (*44*), microbe-to-host ratios were reconstructed and the microbial load of each sample was quantified. Using a count table scaled by the microbial load, we performed ordinations based on Bray-Curtis dissimilarity matrices using R package phyloseq (*72*) and visualised it in a Constrained analysis of principle coordinates (CAP). We performed Permutational Multivariate Analysis of Variance (PERMANOVA) with 999 permutations with the adonis function from the vegan package to quantify the effects of our experiment. To identify taxa differentially abundant across genotypes, microbiomes, and their interaction, we used the DESeq2 (*45*) package (v1.38.3) in R. A full factorial model was fitted with the formula ∼ *line* * *microbiome*, where *line* denotes host genotype (WT and *mmf1-2*) and *microbiome* referred to BX_minus_ and BX_plus_ soils. This design allowed for estimation of main effects and interaction terms while accounting for variance structure and normalization of count data using DESeq2’s internal size factor estimation. ASVs were considered significantly differentially abundant for a given model term if they had an adjusted *p*-value (Benjamini–Hochberg FDR) < 0.05. ASVs significant for any of the three terms (*line*, *microbiome* and their interaction) were retained for downstream visualization. To aid interpretability of the heatmap, group-wise average relative abundances (collapsed by genotype × microbiome) were computed from normalized counts and square-root transformed to reduce the influence of highly abundant ASVs. ASVs were first grouped based on their significance in the DESeq2 model as follows: (1) *microbiome-only*, (2) *line-only*, (3) *interaction-only*, and (4) *multi-category* (significant for multiple terms). Within each group, ASVs were sorted by their average abundance across all samples. Genus-level taxonomy was appended to each ASV ID where available. The microbiome data is in **Dataset 5** and its analysis is documented in **Suppl. File 5**.

### Transcriptome analysis

#### RNA extraction, library preparation and sequencing

Roots from 6-week-old Arabidopsis plants were carefully removed from the soil. Excess soil was removed by briefly rinsing the roots in sterile ddH_2_O and dried using a paper towel. The roots were then flash frozen in liquid nitrogen and homogenized in a 1.5 mL Eppendorf tube using a plastic micro pestle (Carl Roth, Karlsruhe, Germany). Total RNA was extracted using the RNeasy Plant Mini Kit (Qiagen, Hilden, Germany) with on-column DNase treatment with the RNase-Free DNase kit (Qiagen, Hilden, Germany) following the manufacturer’s protocol. RNA integrity was assessed using a Fragment Analyzer (Agilent Technology, Santa Clara, California, USA). RNA concentration was determined fluorometrically using the Qubit RNA HS Assay Kit (Thermo Fisher, Waltham, Massachusetts, USA). Libraries were prepared using the TruSeq Stranded mRNA kit (Illumina, SanDiego, California, USA) with 200 ng total RNA as input. Library preparation and sequencing was performed at the Genomics Facility (https://bsse.ethz.ch/genomicsbasel) of the University of Basel. Sequencing (51 bp paired end) was performed on an Illumina Novaseq® 6000 System (Illumina, SanDiego, California, USA) with an SP slow cell.

#### Bioinformatics and statistics

Reads were aligned to the *Arabidopsis thaliana* TAIR10 reference genome and read counts determined using STAR 2.7.9a (*73*). The raw data processing was performed at the scientific computing facility of the University of Basel (http://scicore.unibas.ch/). Sample outliers were identified using a combination of generalized principal component analysis and by analysing Poisson distances across samples. Sample clustering was performed using variance stabilizing transformations on the normalized gene counts. Differential gene expression analysis was performed in R using DESeq2 (*45*), a method for differential analysis of count data using a negative binomial generalized linear model with shrinkage estimation of dispersions and fold changes. Co-expression analysis was performed with the R package coseq v1.18.0 (*74*). Clustering of the co-expressed genes was performed by running a Gaussian mixture model and arcsine transformation of normalized expression profiles 20 times to identify the optimal number of clusters (*74*). The transcriptomic data is in **Dataset 6** and the documented analysis in **Suppl. File 6**.

### Structural modelling

We utilized the AlphaFold 3 webserver (https://alphafoldserver.com) (*75*) to generate mono- and tetrameric models. Each model was predicted using amino acid sequence of full-length MMF1 with ADP (for monomer model) or with four ATP and four Mg^2+^ (for tetramer model), respectively. The resulting models and RPP1 resistosome structure (PDB: 7CRC) were analyzed using ChimeraX version 1.8.

### Statistical analysis

We used R (R core Team, 2016) for statistical analysis and visualization of data. The R version and package version used for each step of the analysis is documented at the end of the markdown files. All the code required to reproduce the statistical analysis and graphing can be found on the GitHub repository of this article. Figures were exported from R and combined and formatted using Adobe Illustrator. The appropriate statistical test used for each analysis is indicated in the corresponding figure legends and is annotated in the available code.

### Data availability

We have deposited all source data together with all analysis scripts underlying all figures on GitHub (https://github.com/PMI-Basel/JansevanRensburg_et_al_GWAS_MMF1). The GitHub repository’s structure follows the structure of this manuscript with separate folders of source data and analysis scripts on i) Accession growth feedback (Dataset 1), ii) MBOA sensitivity (Dataset 2), iii) GWAS statistics (Dataset 3), iv) Col-0 and mutant growth feedback (Dataset 4), v) microbiome (Dataset 5) and vi) transcriptome (Dataset 6). Each dataset and its analysis documentation (R-markdown outputs) is shared with this article as **Dataset 1** to **6** and the corresponding **Suppl. Files 1** to **6**. The raw sequencing data are stored at the European Nucleotide Archive (http://www.ebi.ac.uk/ena). The microbiome profiling is stored under project number PRJEB85607, the transcriptome data is available under PRJEB85599 and the genome of the *mmf1-2* mutant under PRJEB85789. The bioinformatic code of all analyses, including information on barcodes, primers and sample assignments is provided on GitHub. Raw images are available upon request.

### Code availability

All the code required to reproduce the statistical analysis and graphing can be found on GitHub https://github.com/PMI-Basel/JansevanRensburg_et_al_GWAS_MMF1.

## Acknowledgements

We specifically thank Nicolas Widmer from Agroscope Changins (Switzerland) for field access and assistance. We thank Prof. Jane Parker for providing Arabidopsis mutant seeds. We thank the team at the Plant Sciences Facility at Vienna BioCenter Core Facilities GmbH (VBCF), a member of the Vienna BioCenter (VBC), Austria, for their help with Arabidopsis image acquisition. We are very grateful to Dr. Pamela Nicholson and her team at the Next Generation Sequencing Platform in Bern for excellent technical support. We thank Dr. Phillipe Demougin and Dr. Christian Beisle (both University of Basel, Switzerland) for their support with RNAseq. Finally, we thank Duncan Crosbie (LMU Munich) and Helmut Blum and Stefan Krebs (Gene Center, LMU Munich) for support with genome sequencing of *mmf1-2*.

## Author contributions

H.J.v.R., N.S., C.B., and K.S., designed the research; H.J.v.R. and K. St. performed greenhouse and S.C. the field soil conditionings; K.J and N.S. performed the phenotyping of accessions and GWA analysis; H.J.v.R. performed GWA candidate gene selection, MBOA sensitivity assays, genetic characterisation of Arabidopsis mutants, Arabidopsis mutant feedback, microbiome and transcriptome experiments and analysis; J.W., performed bioinformatic analysis for the microbiome and transcriptome analyses; H.A. performed structural modelling; C.B., provided funding and access to high-throughput phenotyping infrastructure; H.J.v.R. and N.S. analysed the data; H.J.v.R., N.S. and K.S. wrote the manuscript. All authors revised the paper.

---

**Extended Data Fig. 1.**
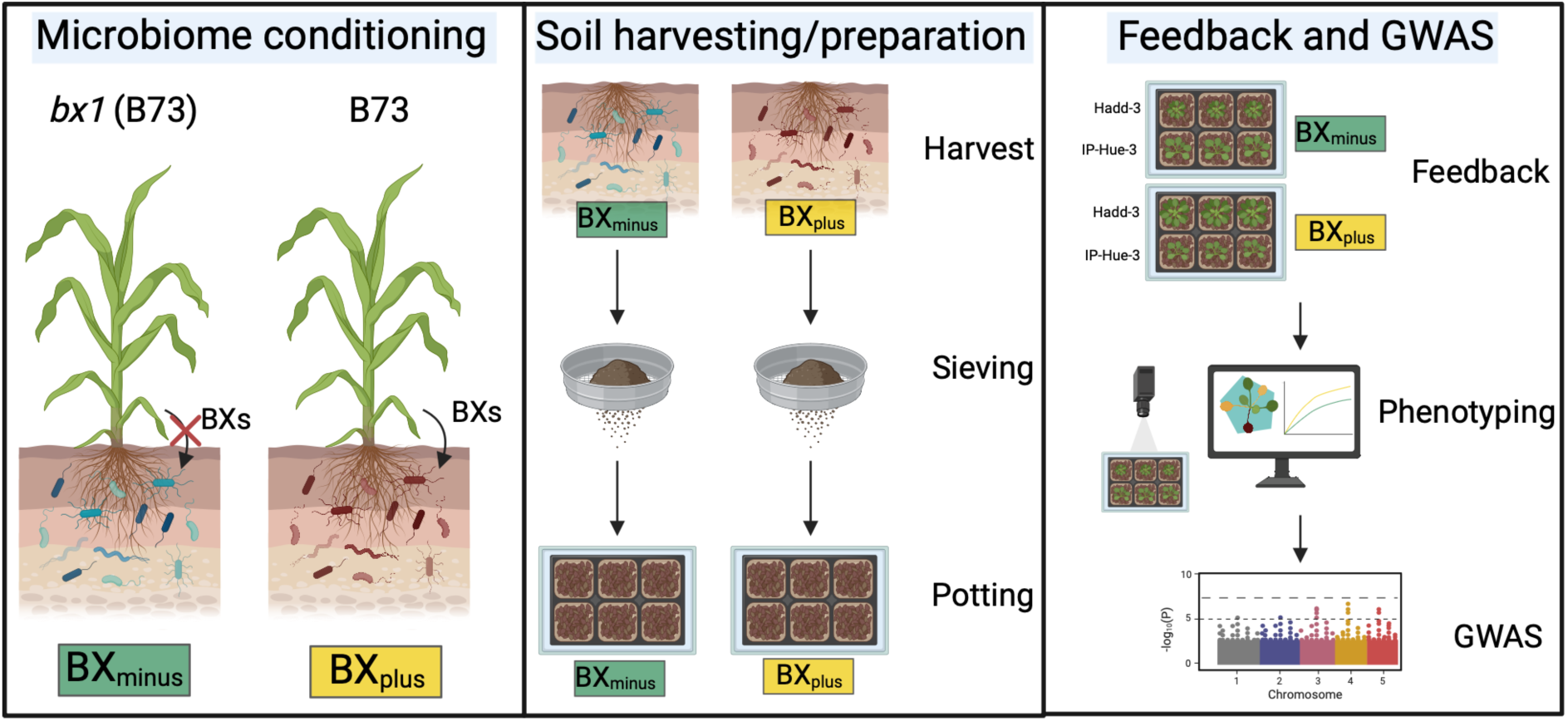
Overview of the experimental workflow. To condition the soil microbiomes, maize exuding BXs (B73) and a BX-deficient mutant (*bx1*) was grown for 12 weeks on field soil, resulting in a BXplus and a BXminus microbiome. The soil core surrounding the roots was removed and sieved to produce the soil used for growth-feedback assays. 410 Arabidopsis accessions were grown on the two conditioned soil microbiomes and phenotyped using a high-throughput phenotyping pipeline. Growth feedback response as the rosette size (BXplus – BXminus) on the two soil microbiomes was used as the phenotype for the genome-wide association study (GWAS). *Mediator of microbiome feedbacks 1* (*MMF1*) identified in the GWAS was functionally validated for growth feedback assays using T-DNA insertion mutant lines. The cause for lack of feedback in mutants were further dissected by analysing the microbiome and transcriptome of wild-type and mutant plants grown on the two microbiomes. Created with BioRender®.

**Extended Data Fig. 2.**
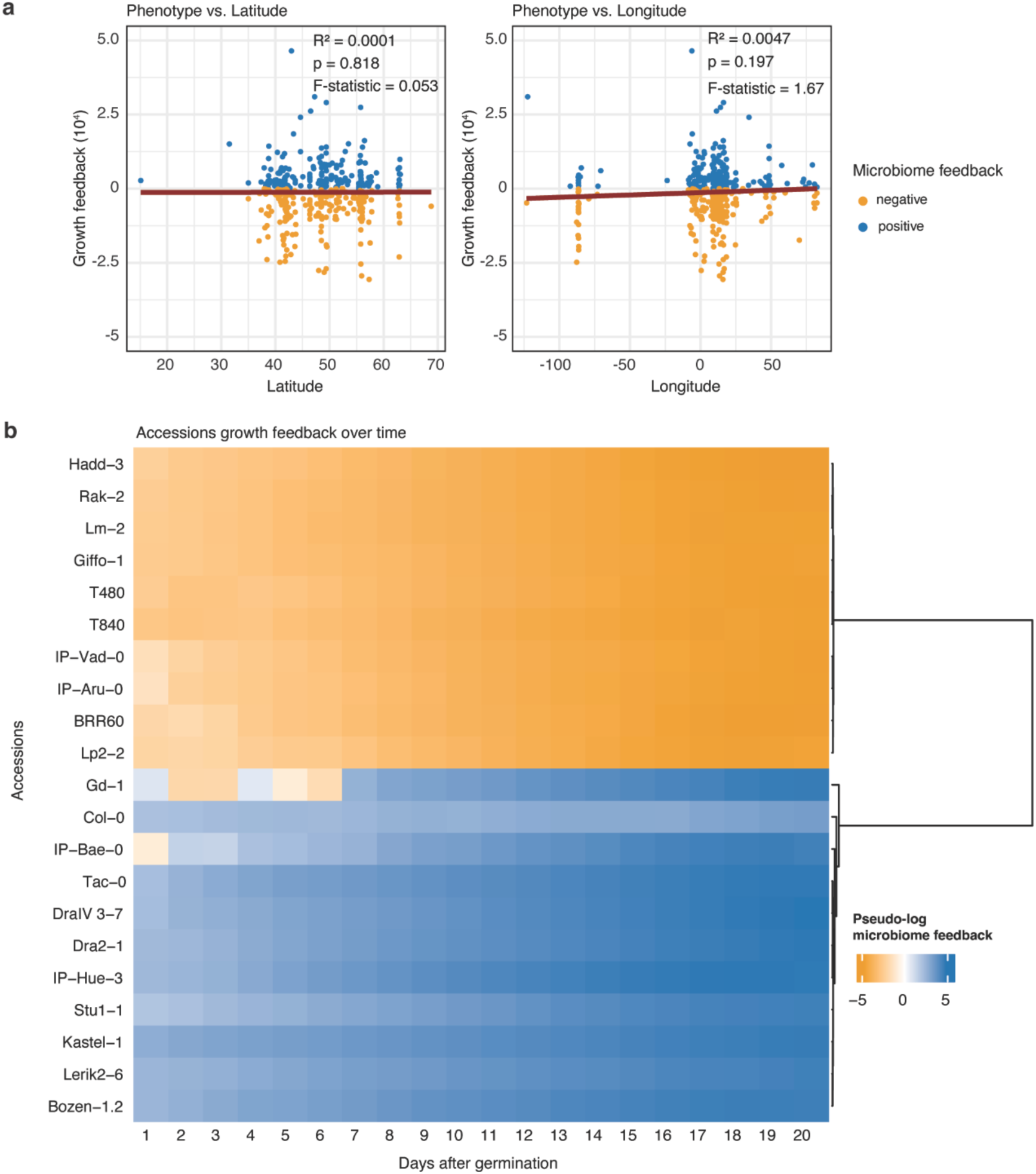
Microbiome feedback of Arabidopsis accessions. (**a**) Relationship between growth feedbacks and geographical location of the phenotyped accessions. Scatter plot illustrating the relationship between latitude or longitude and the microbiome feedback of accessions at 15 days after germination. Each data point is coloured according to its microbiome feedback response, with orange and blue denoting a negative or positive microbiome feedback, respectively. The plot includes a linear regression line (in red), highlighting the relationship between location and phenotype. The regression statistics, including the R-squared value (*R²*) and the *p*-value of the F-test, are displayed in the top right corner of the plot. The R-squared value quantifies the proportion of variance in the phenotype explained by either latitude or longitude, while the *p*-value refers to testing the null hypothesis that the slope of the regression line is zero. (**b**) The daily microbiome feedback of Col-0 and the top 10 accessions with the strongest positive (blue) or negative (orange) BX-growth feedback when grown on the BXplus and BXminus soils. The heatmap colours are the pseudo-log transformed microbiome feedbacks (rosette area on BXplus – rosette area on BXminus). Accessions are clustered on the y-axis based on agglomerative hierarchical clustering using Ward’s method performed using Dynamic Time Warping (DTW) distances and Ward’s method to identify groups with similar response patterns over time. The dendrogram on the right side of the heat map illustrates the clustering structure, grouping accessions that exhibit similar trajectories in their growth feedback **Dataset 1**.

**Extended Data Fig. 3.**
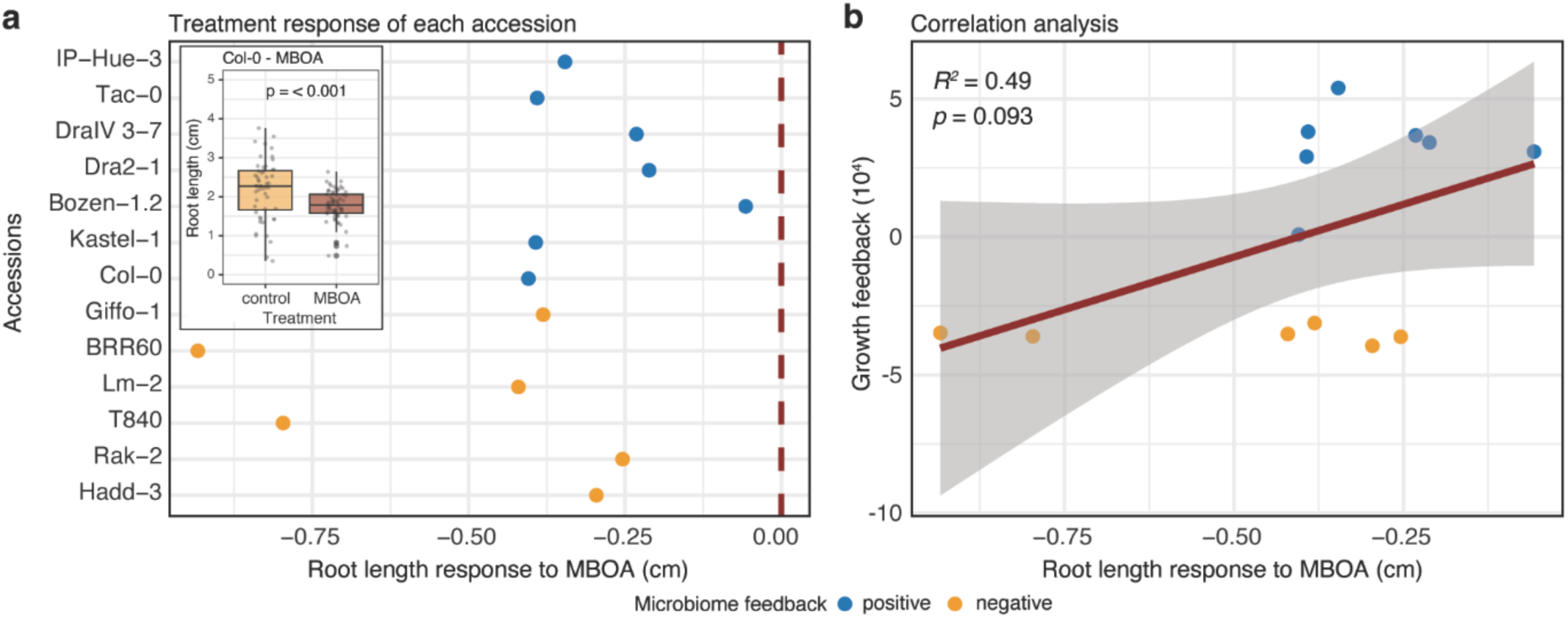
Direct allelopathic effect of benzoxazinoids on the growth of Arabidopsis. (**a**) Sensitivity to MBOA in terms of root length response (root length on control – root length on MBOA) of Col-0 and the accessions with the strongest positive or negative microbiome feedback. Accessions are colour-coded as to whether they showed a positive (blue) or negative (orange) microbiome feedback on the BX-conditioned soil microbiome in the GWAS experiment. Boxplot insert displays the root length of Col-0 on the control or MBOA treatment as an example. Plants were grown for 10 days on ½ MS media in the presence of 0.25 mM MBOA or control (DMSO). (**b**) Pearson’s correlation analysis between MBOA sensitivity and microbiome feedback at 15 days after germination across the different accessions. The scatter plot displays individual data points coloured by microbiome feedback, with positive responses shown in blue and negative responses in orange. A linear regression line (in red) illustrates the trend between the two variables. The correlation coefficient (*R^2^*) and *p*-value are annotated on the plot, indicating the strength and statistical probability of the relationship. **Dataset 2.**

**Extended Data Fig. 4.**
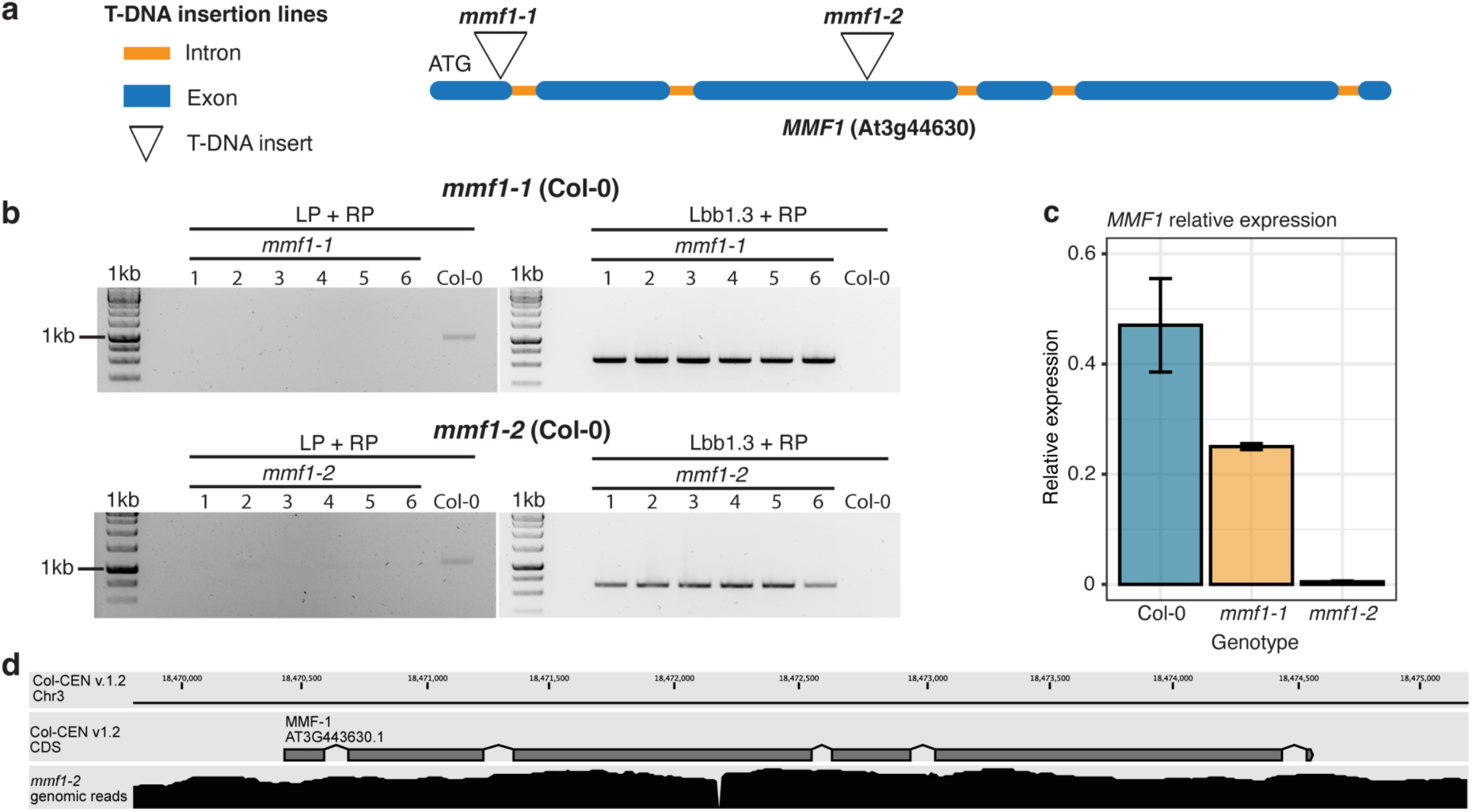
Genetic characterization of *mmf1* T-DNA insertion lines. (**a**) Genomic location of the T-DNA insertion sites for *mmf1-1* and *mmf1-2* in the open reading frame of *MMF1*. Exons and introns are shown in colour, and the T-DNA insertion position as triangles. (**b**) Genotyping results of *mmf1-1* (SALK_144159C) and *mmf1-2* (SALK_002318C) by PCR to confirm the presence of the T-DNA insertion. The gels represent genotyping of 6 mutant plants for the T-DNA insertion allele (left, LBb1.3 and RP primers) and the wild-type allele (right, LP and RP primers). (**c**) Relative expression levels of *MMF1* determined by qRT-PCR in wild-type (Col-0) and the two *MMF1* mutant lines. (**d**) Mapping of genomic long-reads from *mmf1-2* onto Col-CEN v1.2. Top panel shows the genomic coordinates on chromosome 3, middle panel shows the annotation of the coding sequence of *mmf-1,* bottom panel shows the aggregated pile-up of mapped reads, confirming the T-DNA insertion in the third exon.

**Extended Data Fig. 5.**
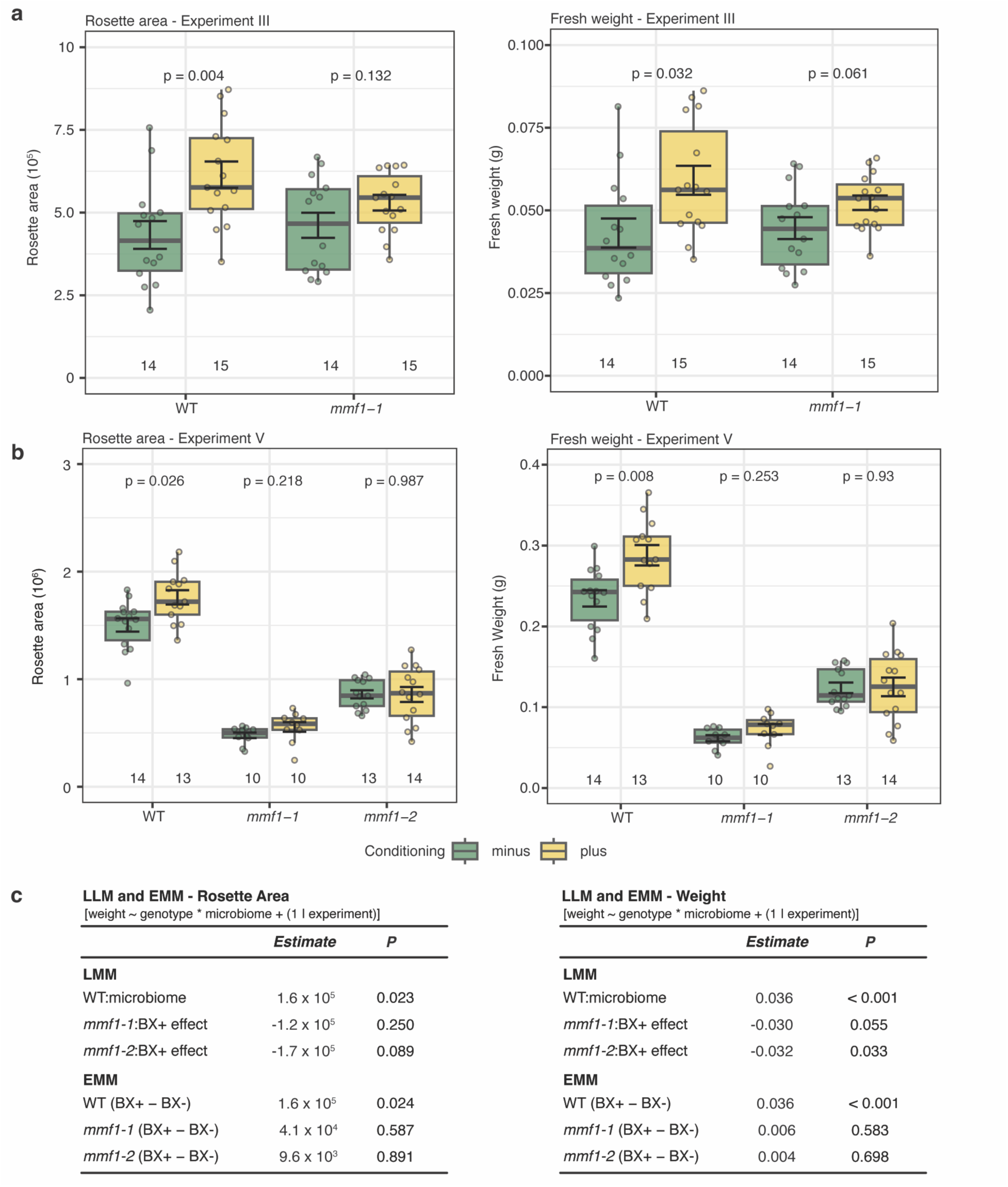
Growth feedback in a BXplus microbiome is consistently lost in *mmf1* mutants. (**a** and **b**) Repeated growth feedbacks (Experiment II and IV) of Col-0 wild-type (WT) and two *mmf1* T-DNA insertion mutants (*At3g44630*) grown on the BXminus and BXplus soil microbiomes. Feedback experiments were performed on greenhouse-conditioned soil batches (**a**) BS04 and (**b**) BS09. Graphs illustrate the rosette area (in pixels) of 6-week-old plants. Sample sizes are indicated at the bottom of each plot. Output of the ANOVA and FDR corrected *p*-values are indicated in each plot. (**c**) A linear mixed-effects model and estimate marginal means (fresh weight/rosette area ∼ line * BX + (1 | experiment)) was used to assess the effects of plant genotype (line), on the growth response to the two microbiomes, with experiment included as a random effect. Data from experiment V and VI were included in the model. **Dataset 4.**

**Extended Data Fig. 6.**
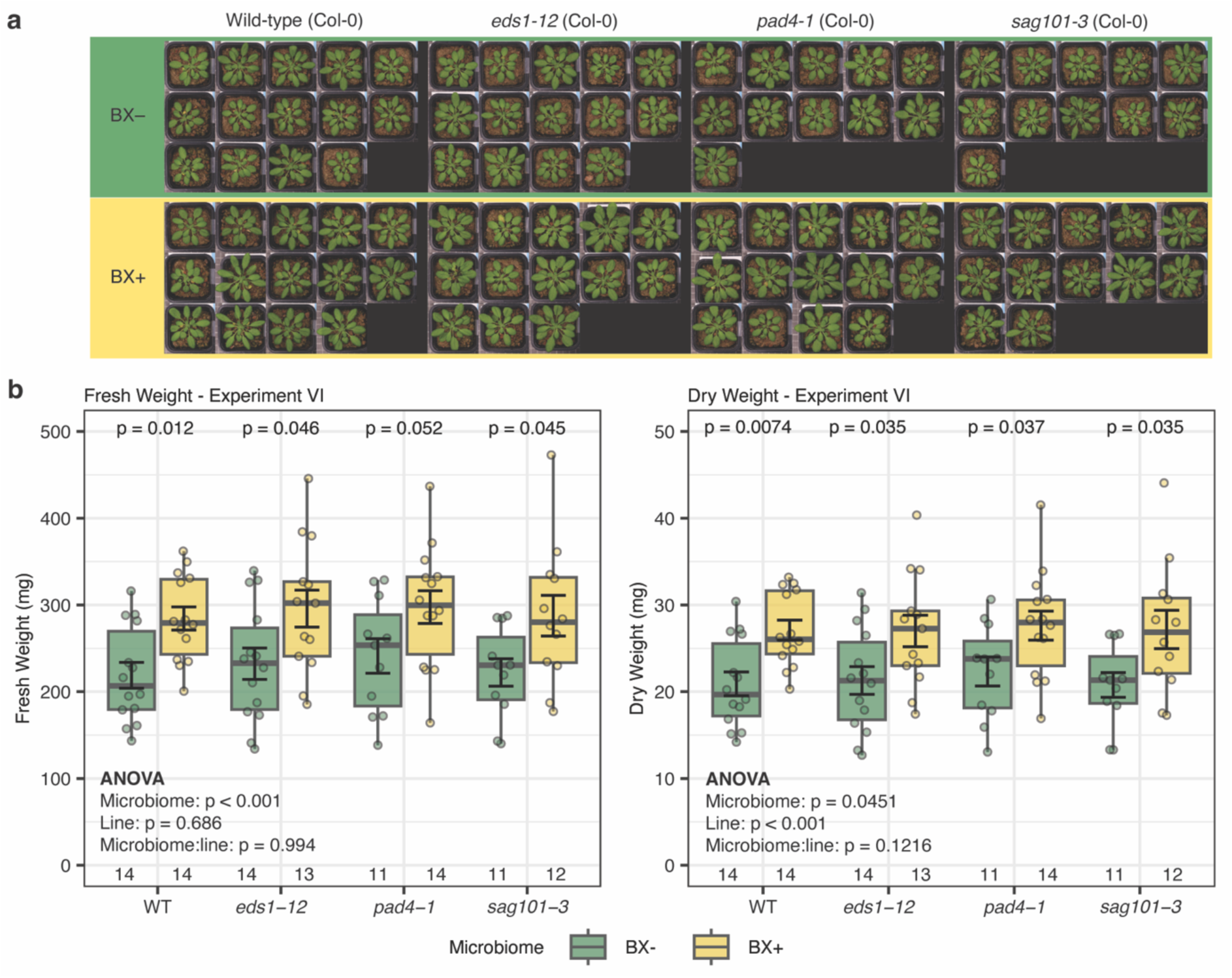
Growth feedback in a BXplus microbiome remains intact in effector-triggered immunity signalling mutants. (**a**) Phenotypes of 6-week-old WT, and mutants (*eds1-12*, *pad4-1* and *sag101-3*) affected in effector-triggered immunity signalling pathways grown on the BXminus and BXplus soils. (**b**) Fresh weight, and dry weight of WT, *eds1-12*, *pad4-1* and *sag101-3* illustrate the positive growth feedback on the BXplus soil microbiome. Boxplots illustrate the median, standard error, and interquartile range, and the whiskers extend from the box to the smallest and largest values. Sample numbers are indicated at the bottom of each plot. ANOVA output and FDR-corrected *p*-values comparing the growth feedback of each line are shown above the graph. **Dataset 4.**

**Extended Data Fig. 7.**
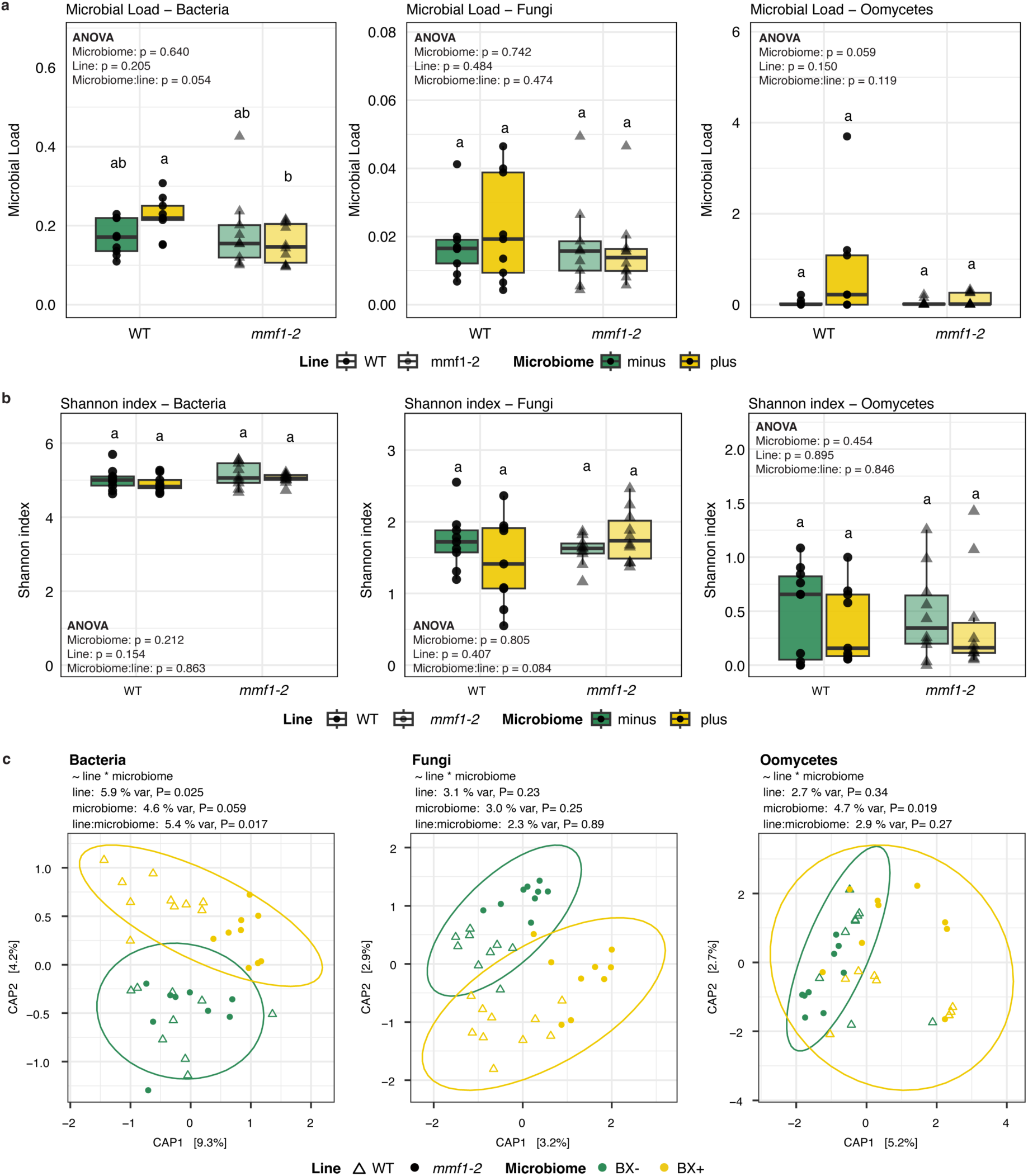
Root microbiota profiling of WT and *mmf1-2* mutants grown on the BXminus or BXminus soils. (**a**) Bacterial, fungal and oomycete load in the roots of wild-type (WT) and mmf1-2 mutants grown on BXplus and BXminus soil. Bacterial load calculated by dividing microbial abundance by host abundance (GIGANTEA). Comparisons between the two lines are shown as *p*-values based on Kruskal-Wallis Test. (**b**) Microbial richness in the roots of the same plants grown on the two soils displayed as Shannon indices. Individual samples are shown as dots. Comparisons between the two lines are shown as *p*-values based on Kruskal-Wallis Test. (**c**) The output of PERMANOVA on Bray-Curtis dissimilarities of the bacterial, fungal and oomycete root microbiota of Arabidopsis WT (filled circles) and *mmf1-2* mutants (open triangles) grown on the BXminus (green) and BXplus (yellow) soils. The *R^2^* and *p* values indicate the line and conditioning effect and their interaction on the two soils. Constrained Analysis of Principal Coordinates (CAP) visually illustrating the effects found in the PERMANOVA analysis. Plants grown on a BXplus (yellow) and BXminus (green) soil microbiome are grouped based on a 95% confidence interval. The axes show the percentage of explained variance for the first two principal components.

**Extended Data Fig. 8.**
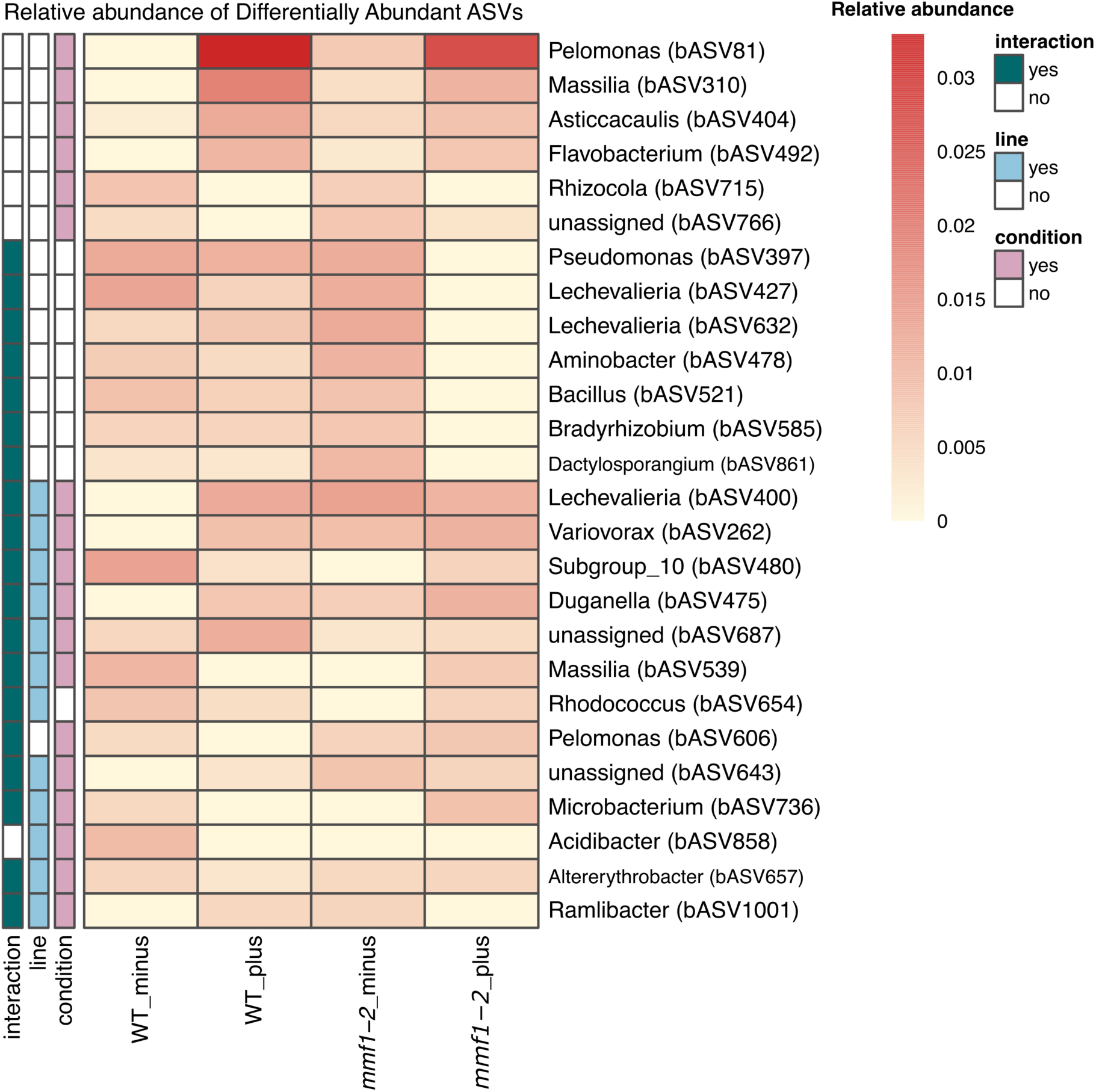
Relative abundances of differentially abundant ASVs in the roots of wild-type and *mmf1* mutants. Square-root-transformed relative abundances of differentially abundant ASVs in the roots of wild-type and *mmf1-2* mutant plants grown on the BXminus or BXplus soil microbiomes. Differential abundance was assessed using DESeq2 with a full factorial model including genotype (*line*), microbiome (*condition*), and their interaction. ASVs were considered significant if they had an absolute log₂ fold change ≥ 1 and a false discovery rate (FDR) < 0.05. Significant ASVs were grouped by the model term(s) they were associated with: condition (pink), genotype (light blue), or interaction (green). Within each group, ASVs are sorted by their average abundance. Genus-level taxonomic annotations are shown, and in parentheses the ASV IDs.

**Extended Data Fig. 9.**
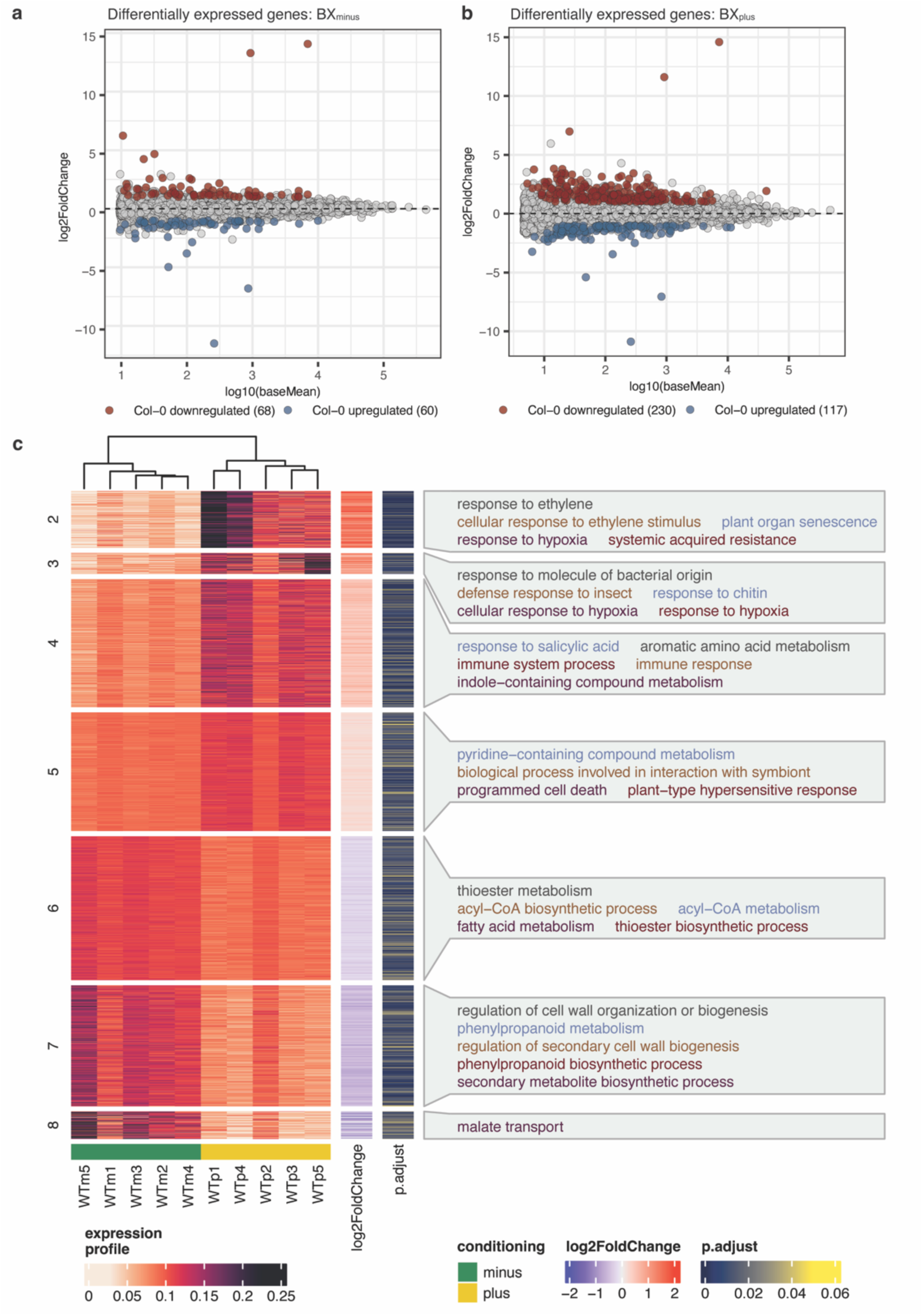
Root transcriptome of WT and *mmf1-2* mutants grown on the BXminus and BXplus soils. MA plot of differentially expressed genes between *mmf1-2* mutants and WT plants in response to the (**a**) BXminus and (**b**) BXplus soil microbiome. Significantly differentially expressed genes were calculated using DESeq2 (log2fold change ≥ 1; empirical Bayes statistic false discovery rate < 0.05). (**c**) Heatmap showing the co-expressed gene clusters (Benjamini-Hochberg-correction, p-adjust < 0.01) as identified by coseq for plants grown on BXminus (green) or BXplus (yellow) soils. Colours of the heatmap indicates the expression profiles in terms of normalized gene counts for each gene. The log2Fold change and adjusted *p* values (empirical Bayes shrinking) for each gene inferred from DESeq2 is shown on the right. Cluster labels show the top 5 enriched GO terms for biological function within each cluster. GO text colour is for better readability only. Full tables of all enriched GO terms in each cluster are provided in **Dataset 6**.

