## Supplementary Materials for "A TNL receptor mediates microbiome feedbacks in Arabidopsis"

**Title:**

**The file includes:**

Supplementary results.

Figures S1 to S2

Tables S1 to S3

**Other Supplementary Materials for this manuscript include**

Dataset S1 to S6 (Input data to reproduce figures)

Supplementary Files 1 to 6 (R markdown files to reproduce analysis)

### Supplementary Results

***Structural modeling of MMF1***

We first assessed the relationship of MMF1 and other TNLs of Arabidopsis using BLAST. MMF1 had highest sequence similarity to the well-characterized sensor TNL Recognition of Peronospora Parasitica 1 RPP1 (*1*). At the protein level, MMF1 shared 84% sequence similarity with RPP1. To investigate the structural organization of MMF1, we generated mono- and tetrameric models using AlphaFold 3 (*2*). The predicted monomer model was generated in the presence of ADP (i.e., in its resting state). The predicted template modelling (pTM) score for the monomer was 0.64, reflecting moderate confidence in domain packing, while the high interface predicted template modeling (ipTM) score (ipTM = 0.95) indicated strong confidence in the relative arrangement of the domains within the monomeric unit (**Fig. S1a**). We noticed high pLDDT values for most domains but not at the terminal extensions. MMF1 displayed the canonical organization of TNL immune receptors comprising TIR (92–259), NB-ARC (260–605), LRR (606–1069) and C-JID (1070–1188) domains (**Fig. S1b**). Unlike other typical TNLs, we noticed however, unusual N-terminal (residues 1–91) and C-terminal extensions (1189–1240).

We next modeled MMF1 as a tetramer to test for eventual resistosome structure (*1*). The tetramer model included four ATP and four $Mg^{2+}$ molecules bound to the nucleotide-binding domains. The predicted global pTM score for the tetramer was 0.67, and the ipTM score is 0.65, indicating moderate confidence in the overall assembly (**Fig. S1c**). Per-residue pLDDT values highlighted particularly the well-ordered core domains, while flexible regions like the terminal extensions showed lower confidence. The four subunits of MMF1 symmetrically arranged with their LRR domains to the outside and forming an inner pore with the NB-ARC and TIR domains in tetrameric assembly (**Fig. S1d**). This MMF1 tetramer model showed structural similarity to the cryo-EM structure of RPP1 (**Fig. S1e**) (*1*). Overall, structural modelling suggests that MMF1 may belong to the sensor-type of TNLs and that it may form oligomeric structures similar to the previously described resistosome complexes.

***Complementation of mmf1-2 partially restores the growth feedback***

To confirm that *MMF1* is the causal factor in the observed growth feedbacks, we generated complementation lines by introducing a functional *MMF1* allele, driven by the *nopaline synthase* promoter (pNOS), into the *mmf1-2* mutant background. We purposefully chose not to use the native MMF1 promotor, due to the presence of another gene (At3g446320) directly upstream of MMF1. From the positive transformants, we selected two lines with the highest *MMF1* expression levels. Although both lines expressed *MMF1* at lower levels than wild-type plants, line 4-4 had higher expression than line 5-1 (**Fig. S2c**). Correspondingly, the growth feedback was fully restored in line 4-4, but only partially in line 5-1. These results indicate that a critical expression threshold of *MMF1* was reached in line 4-4, sufficient to restore the growth feedback to wild-type levels.

### Supplementary Figures

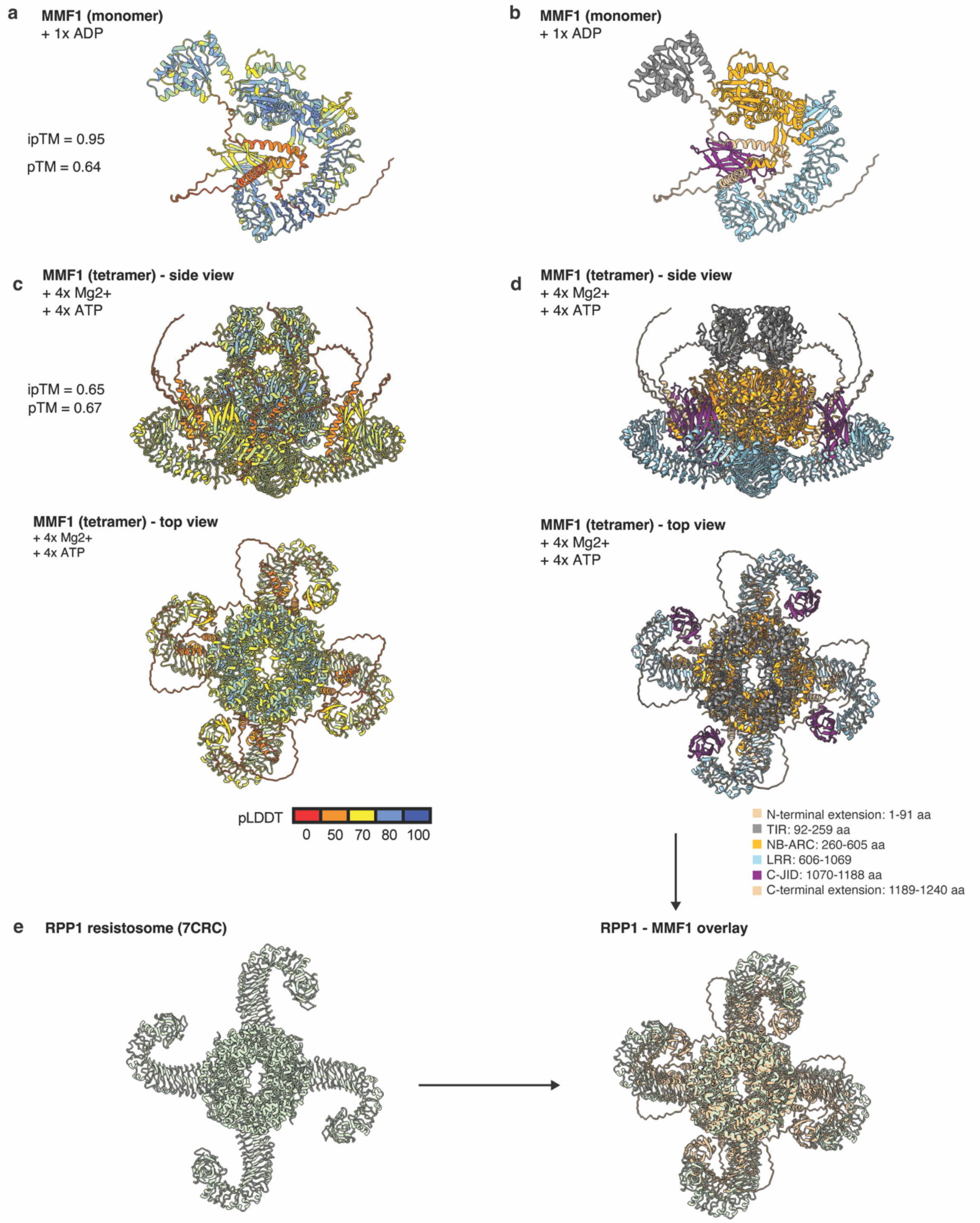


**Figure S1. Structural models of MMF1 based on homology to RPP1.** (**a, b**) Monomeric and (**c, d**) tetrameric models were generated for MMF1 by AlphaFold 3. The template model (TM) confidence scores are indicated as domain packing (ipTM) and global domain arrangement (pTM). Residue-level confidence is shown as a pLDDT-based heatmap (higher pLDDT indicates greater local prediction confidence). Scores **(a)** and domain architecture **(b)** of AlphaFold 3-predicted structure of the MMF1 monomer modeled with ADP. Scores **(c)** and domain architecture **(d)** of tetrameric MMF1 modeled with four $Mg^{2+}$ and ATP molecules. (**e**) Cryo-EM structure of RPP1 and the overlay with the predicted structure of MMF1. Colors legends report the per-residue pLDDT scores and domain structures (incl. segments in amino acids (aa)).

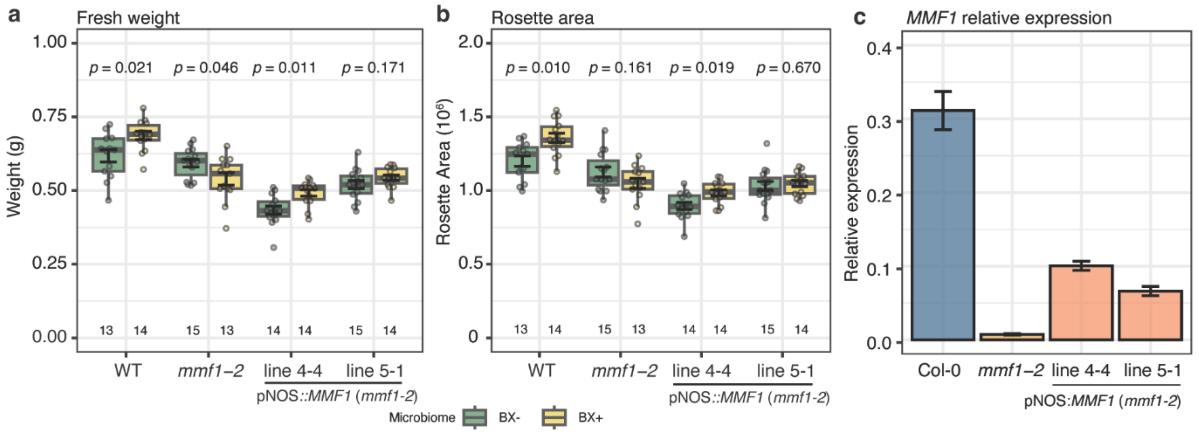


**Figure S2. Growth feedback in a BX$_{plus}$ microbiome is restored *mmf1* complementation lines.** (**a**) Fresh weight and (**b**) rosette area (right; in pixels) of *mmf1-2* mutant lines complimented with *MMF1* under control of the nopaline synthase promotor grown for 6 weeks on the two microbiomes. (**c**) Relative expression levels of *MMF1* determined by qRT-PCR of the two *mmf1-2* lines complemented with *MMF1* under control of the *Nopaline Synthase* Promotor (pNOS).

### Supplementary Tables

**Table S1: Setup and growth conditions of the different experiments performed in this study**. For each of the microbiome feedback experiments in the manuscript, the soil batch, measurement in terms of data generated, and time from start to conclusion is indicated.

| Experiment | Plants | Soil batch | Measurement | Growth (weeks) |
|---|---|---|---|---|
| Exp. I | Accessions run 1 | Vienna1 | Growth (210 accessions) | 12 |
| Exp. II | Accessions run 2 | Vienna2 | Growth (210 accessions) | 9 |
| Exp. III | *mmf1* rep. 1 | BS04 | Growth | 6 |
| Exp. IV | *mmf1* rep. 2 | BS04 | Growth, microbiota, transcriptome | 6 |
| Exp. V | *mmf1* rep. 3 | BS09 | Growth | 6 |
| Exp. VI | ETI mutants | BS07 | Growth | 6 |
| Exp. VII | *mmf1 compl.* | BS10 | Growth | 6 |

**Table S2**: **Set up of the different microbiome conditioning experiments**. Each "soil batch" denotes an independent conditioning experiment from soils collected from three adjacent Agroscope research fields in Changins, Nyon, Switzerland (46°24'00.0"N, 6°14'22.8"E). The "collection" column specifies when and from which field (Parcel) the soil was collected. For fertilization in the greenhouse, pots were treated weekly with the indicated solution. Abbreviations: "rel. hum." = relative humidity; "int." = intensity.

| Soil batch | Collection | Conditioning | Fertilization | Growth conditions |
|---|---|---|---|---|
| Vienna1 | April 2018 (Parcel 30/31) | Field[1] | NA[3] | Conventional farming |
| Vienna2 | Dec. 2019 (Parcel 30/31) | Field[1] | NA[3] | Conventional farming |
| BS04 | Aug. 2020 (Parcel 30/31) | Greenhouse | 4 w. low iron,[4] 8 w. high iron[5] | 16 h day at min. 23°C, 8 h night at min. 19°C, >500 µmol $m^{-2}$ $s^{-1}$ light int. |
| BS07 | Sept. 2022 (Parcel 29/30) | Phytotron[2] | 12 w. low iron[4] | 14 h day at 22°C, 10 h night at 18°C, 60% rel. hum., ~550 µmol $m\text{-}^{2}$ $s^{-1}$ light int. |
| BS09 | Sept. 2022 (Parcel 29/30) | Phytotron[2] | 12 w. low iron[4] | 14 h day at 22°C, 10 h night at 18°C, 60% rel. hum., ~550 µmol $m\text{-}^{2}$ $s^{-1}$ light int. |
| BS10 | Aug. 2023 (Parcel 29) | Phytotron[2] | 12 w. low iron[4] | 14 h day at 22°C, 10 h night at 18°C, 60% rel. hum., ~550 µmol $m\text{-}^{2}$ $s^{-1}$ light int. |

[1]Soil conditioning was performed in the field, and soil cores harvested at the conclusion (collection) of the conditioning.
[2]Phytotron Facility, University of Basel, Switzerland
[3]Fields were managed according to conventional Swiss farming practices, including the use of agrochemicals.
[4]100 mL of 0.2% Plantaaktiv Typ K (Hauert HBG Duenger AG, Grossaffoltern, Switzerland) and 0.001% Sequestrene Rapid (Maag, Westland Schweiz GmbH, Dielsdorf, Switzerland)
[5]200 mL of 0.2% Plantaaktiv Typ K, 0.02% Sequestrene Rapid.

**Table S3: Primer names and sequences used in this study**

| Primer name | Purpose | Primer sequence |
|---|---|---|
| ***mmf1-1*_LP** | Genotyping | GAAGGCTTTTCCAAATTCACC |
| ***mmf1-2*_RP** | Genotyping | ACCAAAATTCCATTAAACCCG |
| ***mmf1-2*_LP** | Genotyping | TAAGGATAATCGGGATTTGGG |
| ***mmf1-2*_RP** | Genotyping | TCCTGAAACATGAGCCAAAAG |
| **LBb1.3** | Genotyping | ATTTTGCCGATTTCGGAAC |
| ***mmf1*_Fw** | *qPCR* | TCGCATGTGAGGCATACGAA |
| ***mmf1*_Rev** | *qPCR* | GCGGTGATGAACGAATTGCT |
| **PP2A_Fw** | *qPCR* | CCTGCGGTAATAACTGCATCT |
| **PP2A_Rev** | *qPCR* | CTTCACTTAGCTCCACCAAGCA |
| **CS1-*fs*-799-F**[1] | Microbiome | ACACTGACGACATGGTTCTACA*fs*AACMGGATTAGATACCCKG |
| **CS2-*fs*-1193R**[1] | Microbiome | TACGGTAGCAGAGACTTGGTCT*fs*ACGTCATCCCCACCTTCC |
| **CS1-At.GI-F** | Microbiome | ACACTGACGACATGGTTCTACACTGTAAAGATAAATGGGTCATCTAA |
| **CS2-At-GI-R** | Microbiome | TACGGTAGCAGAGACTTGGTCTAAGGGTTCAGCTTTGTCAACAA |
| **ITS1-F** | Microbiome | ACACTGACGACATGGTTCTACACTTGGTCATTTAGAGGAAGTAA |
| **ITS2-R** | Microbiome | TACGGTAGCAGAGACTTGGTCTGCTGCGTTCTTCATCGATGC |
| **ITS1-O_F1_G-46636** | Microbiome | ACACTGACGACATGGTTCTACACGGAAGGATCATTACCAC |
| **5.8s-O_R1_G-46637** | Microbiome | TACGGTAGCAGAGACTTGGTCTAGCCTAGACATCCACTGCTG |

[1]*fs* – frame-shift primers to introduce nucleotide heterogeneity for optimized Illumina-based amplicon sequencing. Equimolar pooled set of 5 unique primers.
